# Plants stand still but hide: imperfect and heterogeneous detection is the rule when counting plants

**DOI:** 10.1101/2022.09.05.506614

**Authors:** Jan Perret, Aurélien Besnard, Anne Charpentier, Guillaume Papuga

## Abstract

1. The estimation of population size and its variation across space and time largely relies on counts of individuals, generally carried out within spatial units such as quadrats or sites. Missing individuals during counting (i.e. imperfect detection) results in biased estimates of population size and trends. Imperfect detection has been shown to be the rule in animal studies, and most studies now correct for this bias by estimating detection probability. Yet this correction remains exceptional in plant studies, suggesting that most plant ecologists implicitly assume that all individuals are always detected.
2. To assess if this assumption is valid, we conducted a field experiment to estimate individual detection probability in plant counts conducted in 1×1 m quadrats. We selected 30 herbaceous plant species along a gradient of conspicuousness at 24 sites along a gradient of habitat closure, and asked groups of observers to count individuals in 10 quadrats using three counting methods requiring progressively increasing times to complete (quick count, unlimited count and cell count). In total, 158 participants took part in the experiment, allowing an analysis of the results of 5,024 counts.
3. Over all field sessions, no observer succeeded in detecting all the individuals in the 10 quadrats. The mean detection rate was 0.44 (ranging from 0.11 to 0.82) for the quick count, 0.59 for the unlimited count (range 0.18–0.87) and 0.74 for the cell count (range 0.46-0.94).
4. Detection probability increased with the conspicuousness of the target species and decreased with the density of individuals and habitat closure. The observer’s experience in botany had little effect on detection probability, whereas detection was strongly affected by the time observers spent counting. Yet although the more time-consuming methods increased detection probability, none achieved perfect detection, nor did they reduce the effect on detection probability of the variables we measured.
5. *Synthesis*. Our results show that detection is imperfect and highly heterogeneous when counting plants. To avoid biased estimates when assessing the size, temporal or spatial trends of plant populations, plant ecologists should use methods that estimate the detection probability of individuals rather than relying on raw counts.

## Introduction

Population size and the variability of population size over time and space are of central importance in ecology and conservation. These parameters are required to understand ecosystem dynamics (Ripple & Beschta, 2012; Watson & Estes, 2011), assess population viability (Dennis et al., 1991) and trigger conservation policies and actions (e.g. IUCN, 2019). Population size estimates largely rely on counts of individuals, generally made in spatial units such as quadrats or sites (Elphick, 2008; Yoccoz et al., 2001). Therefore, the accuracy of these counts is crucial, as counting errors would result in biased estimates of population sizes and trends.

When carrying out counts, observers can miss some individuals, in effect counting fewer individuals than were present in the sampling units. This phenomenon of imperfect detection, well known in animal studies, leads to underestimating population sizes (Williams et al., 2002). If detection is imperfect but constant over time and space, raw counts can provide unbiased estimates of population trends or spatial variation (Yoccoz et al., 2001). However, if detection varies over time (e.g. across the sampling season or between years) or space (e.g. between habitats), the raw differences between counts performed at different sampling sessions or in different sampling units are a mixture of differences in abundance and detection (Kéry et al., 2009). For example, a decrease in population size over the years may be hidden as observers improve their ability to detect individuals over time (Kendall et al., 1996). Or one might conclude that there are fewer individuals in closed than in open habitats because they are harder to detect in closed habitats (Buckland et al., 2008). Even differences as small as 4–8% in detection dramatically increase the risk of erroneously concluding that population size varies over time or space (Archaux et al., 2012).

Since the 1970s, many studies on vertebrates have shown that imperfect detection is the rule and that detection probability tends to vary in time and space. Detection probability depends on a number of factors: the observation process (both the observation method and the survey effort), the visibility of the target species, and the characteristics of its habitat (Kéry et al., 2005; Kéry & Schmidt, 2008; Petitot et al., 2014). Consequently, studies including an estimate of detection probability, which corrects for the bias resulting from imperfect detection, have become increasingly common for vertebrates and, to a lesser extent, invertebrates. Yet, they remain exceptional for plants (Kellner & Swihart, 2014), suggesting that most plant ecologists implicitly assume that all individuals are always detected or that differences in detection probability are negligible. It is true that, unlike most animals, plants are immobile, and observers may thus consider it unlikely to miss individuals if careful sampling is conducted. This assumption led John Harper to make the famous statement ‘Plants stand still and wait to be counted’, comparing the ease of studying plant and animal populations (Harper, 1977).

Although detection has almost never been investigated in plants at the individual level, some studies have investigated it at the species level, assessing species occurrence in a given area using two main approaches: occupancy or time-to-detection designs. In the first approach, the survey duration is fixed, and the detection probability of the species when surveying quadrats or sites is estimated from repeated surveys. In the second approach, the detection probability is estimated based on the survey time required to detect the species. These studies found that the detection of plant species was almost always imperfect and that detection probability varied depending on ecological and observational variables. For example, species detection varied according to the morphology of the target species (Chen et al., 2009, 2013; Garrard et al., 2013), with conspicuous growth forms and phenological stages raising detection probability. A high density of individuals increased the probability of detecting the species (Bornand et al., 2014), while the closure of the surrounding habitat decreased it (Alexander et al., 2012; Garrard et al., 2008). Species detection probability also increased with survey effort (Chen et al., 2009) and often varied between observers (Archaux et al., 2006; Garrard et al., 2008. Although these studies confirm that detection issues exist when assessing the occurrence of plant species, do these issues affect individual detection to a similar extent?

Concerning detection at the individual level, the only studies that have been conducted on plants used marked individuals in so-called ‘capture–recapture’ designs (Lebreton et al., 1992) or the time-to-detection of individuals (Hauser et al., 2022). These studies found that individual detection was imperfect, although individuals were generally searched for in relatively small, intensively surveyed plots (Alexander et al., 1997; Andrieu et al., 2017; Kéry & Gregg, 2003). In most studies using marked individuals, individual detection probability varied with the phenological stage of individuals and sometimes with habitat closure (Moore et al., 2011; Andrieu et al., 2017). Hauser et al. (2022) showed that flowering individuals were detected faster than nonflowering ones, and that the abundance of individuals of species resembling the target species delayed detection. Observers with recent experience in searching for the target species detected it faster, confirming at the individual level what has been demonstrated at the species level. Therefore, it is of particular concern that individual detection probability has never been estimated for counts of unmarked plants, despite this being one of the most commonly used methods for estimating population sizes and trends in plant ecology (Elzinga et al., 1998; Reisch et al., 2018). In addition, many studies in plant ecology estimate vital rates by counting unmarked individuals per life stage (e.g. Molano-Flores & Bell, 2012; Giljohann et al., 2017). In theory, data from capture-recapture designs could be used to estimate detection probability and correct estimates from counts of unmarked individuals. To our knowledge, this has never been done, probably because it would be very impractical since both observation processes would have to be strictly equivalent regarding detection and because estimating detection probability directly from counts of unmarked plants is usually less labour–intensive (Royle, 2004).

To assess whether imperfect detection of individuals is indeed the rule in plant counts, we conducted a field experiment in which observers counted plants in 1×1 m quadrats, a commonly used quadrat size (Elzinga et al., 1998; Gauthier et al., 2017, 2019). The experiment covered 30 species and 300 quadrats, and involved 158 observers who made 5,024 counts. Our study had three aims: (i) to estimate the detection probability of individuals in plant counts; (ii) to estimate how detection probability varies in time and space; (iii) to determine if increasing the survey effort results in a higher detection probability, and could allow reaching perfect detection. To meet these objectives, we tested whether detection varies according to ecological variables (species conspicuousness, true population density and surrounding habitat closure) and observational variables (counting method and experience of observers), which are likely to vary over time and space, as well as testing varying degrees of survey effort.

## Materials & methods

### 1 Selection of sites, species and observers

We selected 30 herbaceous plant species from 24 sites throughout France in Mediterranean, Alpine and temperate climatic regions (Appendix 1). We chose species with various growth forms and phenological stages to cover a gradient from cryptic to conspicuous species. As the experiment focused on counting individuals, we selected only species for which ‘individuals’ (ramets or genets depending on the species) could be unambiguously differentiated. We avoided species difficult to identify so that all could be identified at first glance without having to check identification criteria on each detected individual. We selected study sites to cover a gradient of habitat closure, from bare ground to dense meadows, to include all possible combinations of species conspicuousness and habitat closure (Fig. 1).

**Figure 1:**
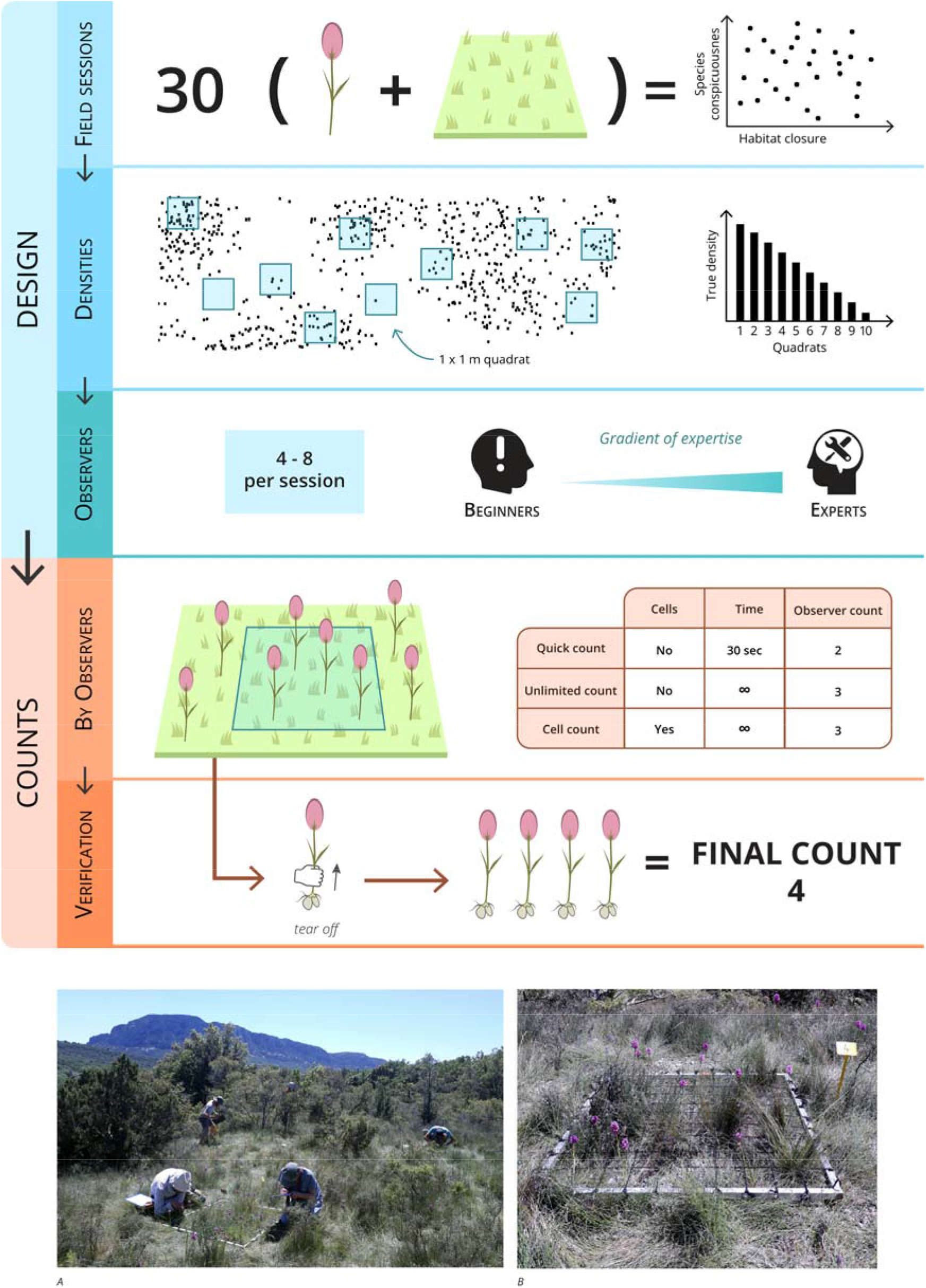
Diagram of the study’s experimental design. We conducted 30 field sessions, each focusing on a different species in a habitat with a different closure level, to cover the widest possible range of combinations of species conspicuousness and habitat closure. For each field session, we placed ten quadrats on the population to form a density gradient and chose 4–8 observers to have a gradient of experience in botany. The observers carried out three counting methods successively (quick, unlimited and cell counts). We then counted the true number of individuals by systematically digging out the individuals detected by the observers and then conducting an additional survey for any individuals that remained unnoticed. Picture A shows the experiment in progress at a Mediterranean site and picture B shows a quadrat with the cells.

Participants were volunteers selected among our colleagues, researchers, undergraduate students and professionals involved in flora conservation. We selected 4 to 8 observers for each field session, including at least one with several years of field experience in botany, one with no experience in botany, and intermediate profiles. Observers were allowed to participate in up to three sessions, but these sessions had to be several weeks apart. This resulted in 171 participations by 158 unique observers. We conducted the fieldwork from 7 October 2020 to 6 August 2021.

### 2 Experiment

#### Field setup and observer training

The fieldwork consisted of one-day sessions, during which a group of observers counted the number of individuals of a single target species in a habitat at one of the 24 sites. The availability of participants restricted us to organise the field sessions. Whenever we had enough observers available somewhere, we prospected the landscape nearby to find an appropriate site (i.e. easy to access and with a habitat corresponding to what we needed to complete the habitat closure gradient). Then we searched for a suitable species, meeting our general criteria and completing the species conspicuousness gradient. Before the observers’ arrival, we placed ten 1×1 m quadrats on the target species population. We chose quadrat locations to ensure that (i) two were free of individuals of the target species (so that observers would not always expect to find individuals in the quadrats), and (ii) they covered a relatively linear gradient of densities up to a maximum of 150 individuals (when possible).

It happened regularly that quadrats meant to be empty contained a few individuals and those with the highest densities contained more than 150 individuals. We labelled the quadrats from 1 to 10 and fixed them to the ground. When the observers arrived, we explained how the counts would be conducted and how to identify the target species at all phenological stages and differentiate it from the other species present at the study site. We then trained them to count on additional quadrats to check that everyone had understood the instructions. This training phase lasted until all observers felt comfortable with species identification and the counting methods. The entire training process usually took about an hour.

#### Counting methods

Each observer had to count the number of individuals of the target plant in each quadrat using three different methods, requiring a progressively increasing amount of survey time. In the first round, participants had 30 seconds to estimate the number of individuals in each quadrat (the quick count). In the second round, they had to count the number of individuals without any time limit (the unlimited count). We then divided the quadrats into 100 cells of 10×10 cm (Fig. 1B), and observers performed the third round, where they had to count the number of individuals in each cell without any time limit (the cell count). This last method forced observers to survey the quadrat homogeneously and has been regularly used in the plant survey literature (e.g. Gauthier et al., 2017, 2019). For the unlimited and cell counts, observers recorded the time they spent counting in each quadrat. To avoid memory effects, after each round we collected the sheets on which the observers recorded their measurements, and observers surveyed the quadrats in a different order, approaching it from a different side at each round. The observers conducted the three counts on the ten quadrats in 1.5 to 5 hours (on average 3.25 hours).

#### Exhaustive count and measurement of covariates

After the observers had left, we carried out a final count to determine the true number of individuals present in each quadrat. This was done by searching each 10×10 cm cell for the maximum number of individuals detected by the observers during the third round and collecting them. Once the maximum number of individuals detected by the observers in each cell had been removed, we searched again for any potentially undetected individuals. This final count took between 1 and 7 hours (3 hours on average), depending on habitat closure and the number of individuals of the target species in the quadrats. Despite our verification method, we might have missed a few individuals during the final count, but we will refer to this as the ‘true number of individuals’ for the sake of simplicity. We calculated the detection probability of individuals by dividing the number of individuals detected by the observers by the true number of individuals present in each quadrat.

We measured three variables to investigate their effect on detection probability: an index of species conspicuousness (measured for each field session), an index of habitat closure (measured for each quadrat), and the experience in botany of the observers. For simplicity’s sake, we used synthetic indices to characterise species conspicuousness and habitat closure. This allowed us to limit the number of variables and interactions in the model (e.g. species conspicuousness combines the average size of individuals, the presence of vividly coloured flowers, and the interaction of these variables). All the variables and interactions that we investigated in the following analysis are presented in Table 1, as well as the underlying hypotheses. The way we measured the variables is detailed in Appendix 2.

### 3 Statistical analysis

In total, 300 quadrats were surveyed during the experiment by on average 5.7 observers, resulting in 5,024 counts. We removed 34 quadrats from the analysis that contained no individuals of the target species, as these do not provide information on detection probability. We withdrew four additional quadrats that unexpectedly contained more than 300 individuals to avoid the leverage effect due to these density outliers (Appendix 3). Keeping these quadrats in the dataset would have led to a reduction in the slope of the density effect, which could result from a nonlinear relationship at high densities, but with only four quadrats, we could not reliably test this hypothesis. In some cases (N = 140, representing 2.8% of the counts), the observers recorded more individuals than were truly present (Appendix 4). We thus carried out the analyses on two datasets, one in which we removed these counts and one in which we set these counts to the true number of individuals. As there was no discernible difference in the results, we present the results of the former analysis in the main text and the latter in Appendix 7. After all filtering steps, 4,319 counts remained in the dataset from which we had removed the observations with excess detections. We also performed the analyses keeping only the experienced botanists to verify that our results were not driven by an excess of novice observers in our sample. Despite some minor differences, our overall results remained unchanged (Appendix 8).

We used a generalised linear mixed model (GLMMs) with a binomial distribution and a logit link function to analyse the correlation between detection probability and the variables we measured and some of their interactions. There could be dependence between detections due to individuals forming clusters. However, observers often detected some but not all individuals within clusters. Furthermore, the aggregation of individuals was not very strong at the quadrat scale. Hence, the dependence between detections is probably relatively small, and we consider the binomial model appropriate. We designed the experiment to disentangle the effects on detection probability of species conspicuousness, habitat closure and survey effort. We had precise hypotheses about the mechanisms explaining these effects (detailed in Table 1) that included interactions between variables, and we collected a large dataset to test all our hypotheses. Therefore, we fitted a single model including all the variables and interactions corresponding to our hypotheses and used their statistical significance for inference. To take into account the repeated measurements at each quadrat, we included random intercepts for the observers, the field sessions and the quadrats (nested within the field session). As the majority of observers participated in only one field session, we did not add a random effect interaction between observers and field sessions. Furthermore, we did not add random effects on the slopes because we had no specific hypothesis that would justify it. We standardised all the variables prior to model fitting and assessed the model’s goodness-of-fit using the package DHARMa (Hartig, 2022). We computed the proportion of variance explained by the model and the proportion of variance explained by the model’s fixed effects, using the theoretical variance method of Nakagawa et al. (2017). Analyses were performed in R version 3.6.1 (R Core Team, 2019), using packages *lme4* (Bates et al., 2022) to fit the model and *ggeffects* (Lüdecke et al., 2022) to plot the predictions.

**Table 1:** Ecological variables (species conspicuousness, habitat closure, true density), observational variables (counting method, experience in botany, quadrat order, counting time), and interactions between variables tested for their effect on the detection probability of individuals, with our hypothesis about the underlying mechanisms. The measurement method of all variables is detailed in Appendix 2.

| Variable | Description | Measurement method | Hypothesis | Type | Observed values |
| --- | --- | --- | --- | --- | --- |
| Species conspicuousness | Conspicuousness of the species at the time of the field session | Expert judgement based on the mean height of individuals and the presence of colourful flowers | Tall and colourful species are easier to detect, which increases their detection probability | Ordinal | 1, 2, 3, 4 |
| Habitat closure | Closure of the vegetation in the quadrat | Composite variable created from the vegetation height and the proportion of the area covered by vegetation in the quadrats (Appendix 2) | Dense vegetation masks individuals, which lowers detection probability | Continuous | -1.86–3.54 |
| True density | Number of individuals present in the quadrat | Number of individuals counted in the quadrat by the experimenter at the end of the experiment | High density requires greater concentration and makes it difficult to distinguish between individuals, reducing detection probability | Count | 0–295 |
| Counting method | Counting method used | Fixed by the experimental design, each observer successively used the three counting methods on the ten quadrats | Counting methods requiring more survey time yield higher detection probability | Categorical | Quick count, unlimited count, cell count |
| Experience in botany | General experience in botany of the observer (not specific to the target species) | Self-evaluated by the observers before the experiment using a standardised form | Observers with more experience in botany have higher detection probability | Ordinal | 1, 2, 3, 4, 5 |
| Quadrat order | Position of the quadrat in the round | Defined by the experimental design, the order in which the observers investigated the quadrats was fixed in advance and different for each observer and round. | Learning increases detection probability over time, or fatigue decreases it | Continuous | 1, 2, 3, ..., 10 |
| Counting time | Time spent counting by the observer | Each observer timed the duration of counting | Spending more time increases detection probability | Continuous | Quick count: 0.5 min<br>Unlimited count: 0–11 min<br>Cell count: 0–41.58 min |
| Species conspicuousness x Habitat closure |  |  | Detection probability is less affected by habitat closure for conspicuous species | Interaction term |  |
| Species conspicuousness x Counting method |  |  | Detection probability obtained with counting methods requiring more survey effort is less affected by species conspicuousness | Interaction term |  |
| Habitat closure x Counting method |  |  | Detection probability obtained with counting methods requiring more survey time is less affected by habitat closure | Interaction term |  |
| Experience in botany x Counting method |  |  | The difference in detection probability between experts and beginners is larger for the quick count than for the other two methods | Interaction term |  |
| True density x<br>Counting<br>method |  |  | Counting methods<br>requiring more<br>survey time make<br>it easier to<br>distinguish<br>between<br>individuals, which<br>decreases the<br>negative effect of<br>true density | Interaction<br>term |  |
| Quadrat order<br>x Counting<br>method |  |  | The effect of<br>fatigue or learning<br>on detection<br>probability is<br>stronger for<br>counting methods<br>requiring more<br>survey time | Interaction<br>term |  |
| Species<br>conspicuousn<br>ess x Counting<br>time x<br>Counting<br>method |  | No coefficient is estimated<br>for the quick count as the<br>time was fixed | Spending an<br>additional extra<br>minute further<br>increases detection<br>probability for<br>species that are<br>easy to detect | Interaction<br>term |  |
| Habitat<br>closure x<br>Counting time<br>x Counting<br>method |  | No coefficient is estimated<br>for the quick count as the<br>time was fixed | Spending an<br>additional extra<br>minute further<br>increases detection<br>probability in<br>sparse vegetation | Interaction<br>term |  |

## Results

Over the 30 field sessions, no observer succeeded in detecting all the individuals in all the quadrats. Averaged over all quadrats and observers, the mean detection rate per field session was 0.44 (ranging from 0.11 to 0.82) for the quick count, 0.59 for the unlimited count (from 0.18 to 0.87), and 0.74 for the cell count (from 0.46 to 0.94).

The goodness-of-fit of the fitted model was very good (Appendix 5). All ecological and observational variables were significantly correlated with detection probability in at least some conditions, i.e. for some counting methods or some levels of the other variables (Fig. 2; Appendix 6). Regarding the ecological variables, as expected, the more closed the habitat, the lower the detection probability (Fig. 3). The more conspicuous the species, the higher the detection probability, except in entirely open habitats, where the detection probability was constant and high regardless of the conspicuousness of the species (Fig. 3). Detection probability decreased steeply with the true density, in a similar way for the three counting methods (Fig. 4). Regarding the observational variables, spending more time sharply increased detection probability for the unlimited and cell counts. Habitat closure moderated this trend, i.e. spending an extra minute counting increased detection probability less in closed habitats than in open habitats. Experienced observers achieved slightly higher detection probability than inexperienced observers, and this difference was stronger for the cell count than for the unlimited count. In contrast, there was no significant correlation between experience and detection probability for the quick count. For the unlimited and cell counts, detection probability slightly increased with the position of the quadrat in the round, while there was no correlation for the quick count (Fig. 4).

**Figure 2:**
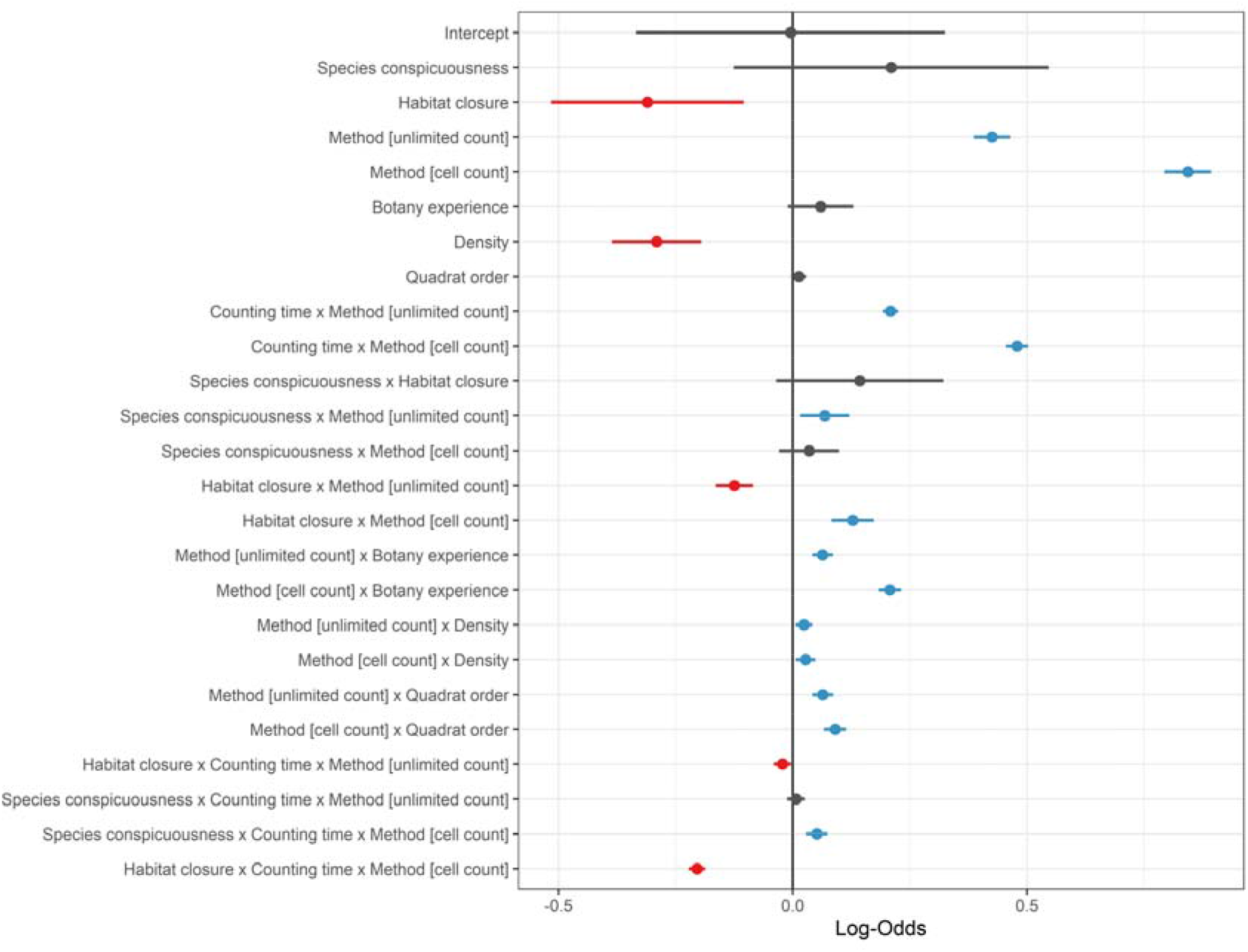
Coefficients of standardised variables in the model. Dots represent estimated coefficients, and lines represent their 95% confidence intervals. Coefficients with a nonsignificant effect are shown in grey. The reference counting method is the quick count, in which observers had 30 seconds to estimate the number of individuals in the quadrat. The coefficients presented for the other two methods indicate the difference (on the logit scale) in mean detection probability obtained with those methods compared to the quick count. The variable ‘counting time’ appears twice because we manually coded the interaction between the counting time and the counting method, so no slope parameter is estimated for the quick count.

**Figure 3:**
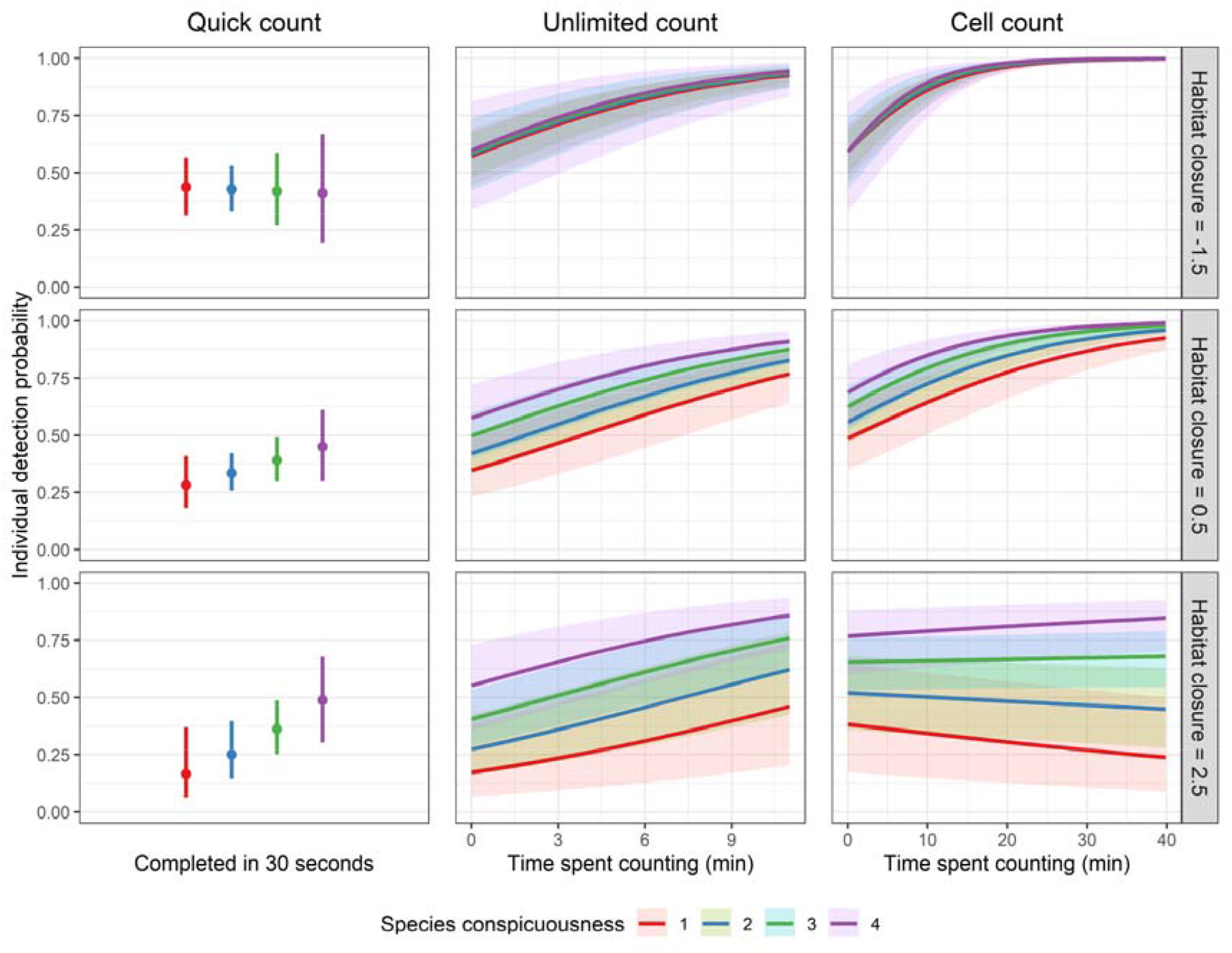
Predicted detection probability depending on species conspicuousness (1 = least conspicuous; 4 = most conspicuous), habitat closure (−1.5 = most open habitats; 2.5 = most closed habitats), the counting method, and the time spent counting. Predictions were made for a quadrat containing 100 individuals of the species of interest, located in the fifth position in the round for an observer who rated his level of experience in botany at 2.5 out of 5. The maximum counting time shown for the unlimited count (11 min) and the cell count (39 min) is the maximum time an observer spent counting on a single quadrat during the experiment. Lines represent the predicted mean detection probability and the shading represents 95% confidence intervals.

**Figure 4:**
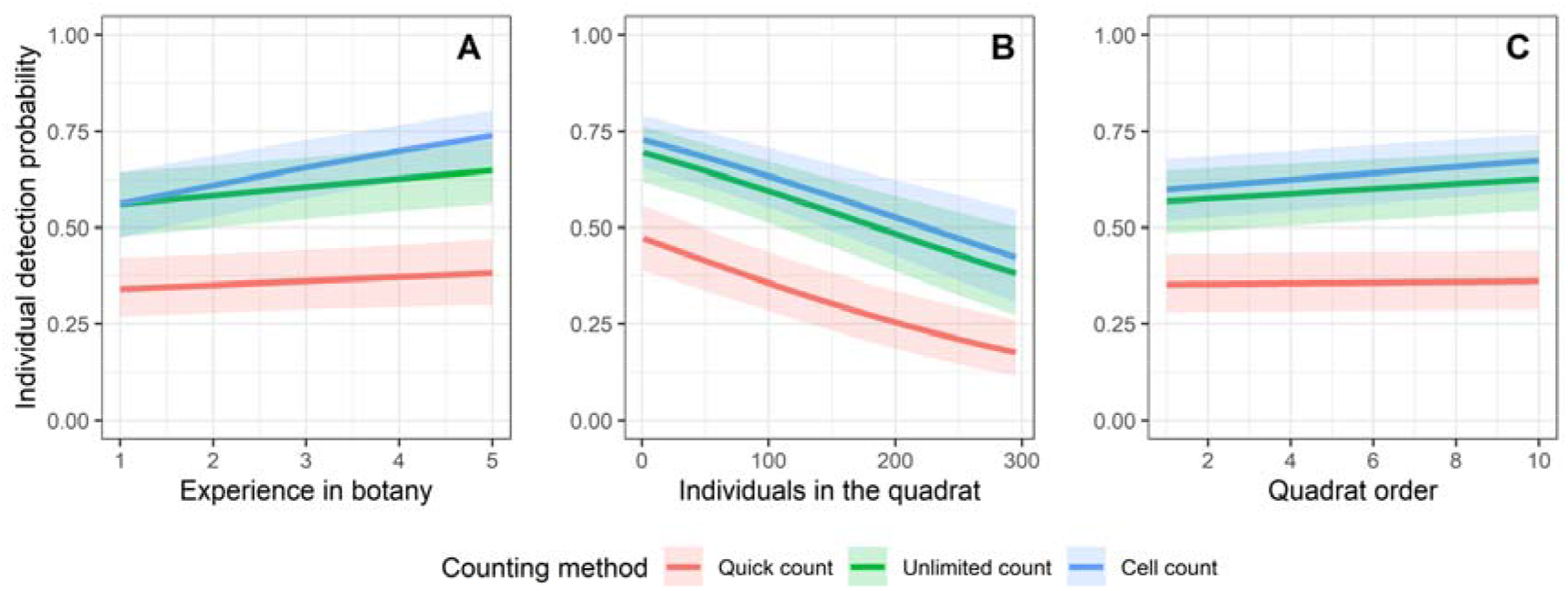
Predicted detection probability of individuals, depending on the experience in botany of the observer (A), the true density (B) and the position of the quadrat in the round (C). The other variables included in the model are held constant at the following levels: species conspicuousness = 2, habitat closure = 0, quick count time = 0.5 min, unlimited count time = 3 min, cell count time = 3 min. Lines represent the mean detection probability of individuals and the shading represents 95% confidence intervals.

Observers achieved higher detection probabilities with the cell count than with the unlimited count, which yielded higher detection probabilities than with the quick count. However, this came at the cost of a major increase in the time spent counting. The counting time per quadrat was fixed at 30 seconds for the quick count, and the mean counting time averaged over all sessions and observers was 1.8 minutes (ranging from 0.3 to 5.7) for the unlimited count and 8.2 minutes (from 0.9 to 21.6) for the cell count. While the time spent increased, the slope between detection probability and the other variables (e.g. observer experience, vegetation closure) was not reduced. Some variables had roughly the same correlation with detection probability regardless of the counting method (e.g. true density), while others had an even steeper slope (e.g. experience in botany).

Within the limits of the counting times we observed during the experiment, the model predicts that reaching a detection probability over 0.95 (i.e. ‘almost perfect detection’) is only possible by using the cell count for quadrats with a true density below 100 individuals and a counting time per quadrat of at least 20 minutes. In a very open habitat (typically with <7% vegetation cover and median vegetation height <1 cm), this can be achieved no matter the species’ conspicuousness. In a less open habitat (e.g. vegetation cover <90%, median vegetation height <6 cm), the target species must be very conspicuous, i.e. a plant above 20 cm in height or 10 cm but with colourful flowers.

Substantial variation in detection probability between field sessions remained unexplained by the model, even after controlling for species and habitat characteristics, as the variance of this random effect was σ_*session*_^2^ = 0.58. There was also unexplained variation between quadrats (σ_*quadray*_^2^ = 0.42). The unexplained between-observer variation was the smallest, with a variance of σ_*observer*_^2^ = 0.16. The proportion of variance explained by the model was *R*^2^_*GLMM*(*m*)_ = 0.35, and the proportion of variance explained by the model’s fixed effects was *R*^2^_*GLMM*(*m*)_ = 0.12. In other words, detection probability remained heterogeneous even after considering the effect of the seven variables included in the model and accounting for unexplained inter-session, inter-plot and inter-observer variance.

## Discussion

This study provides the first estimate of the detection probability of individuals in plant counts, using a detection experiment with a cross-taxa and cross-habitat design. Our results demonstrated that imperfect detection is the rule when counting plants, even in small (1 m^2^) quadrats with unlimited counting time. Detection probability was also highly heterogeneous and varied with the species, habitat, observer and counting method. Methods involving longer survey time increased the mean detection probability, but did not achieve perfect detection, nor did they reduce the effect of the ecological and observational variables on detection probability. Even after controlling for the effects of all the variables, detection remained highly heterogeneous. Therefore, we argue that plant ecologists should not rely on the use of raw counts and should always take into account detection issues when studying plant populations.

### 1. Evidence for imperfect detection of plants

In our experiment, while using common plant counting methods (Reisch et al., 2019), not a single observer detected all the individuals present in all the quadrats of a field session. Even with counts unlimited in time, the predicted detection probability under average conditions (i.e. for the average values of all variables included in the model with the unlimited count) did not exceed 0.59 (95% confidence interval: 0.50–0.67). Thus, our results not only confirm that imperfect detection is the rule when counting plant individuals, but also show that detection probability can even be relatively low. Although these values may be surprising, detection issues have been previously documented for plants when searching for a species (e.g. Chen et al., 2013), and a recent study conducted on large quadrats (20×20 m) suggested that imperfect detection may also occur for plant individuals (Hauser et al., 2022). Furthermore, the detection probabilities we found for plants are higher than those commonly found for animals. For example, detection probabilities of 0.04–0.32 have been found in bird point counts (Royle, 2004), 0.07–0.41 for lizards counted along transects (Kéry et al., 2009), 0.06–0.41 for salamanders counted along stream banks (Dodd & Dorazio, 2004), and 0.06–0.34 for freshwater mussels searched on riverbeds (Carey et al., 2019). Nevertheless, this does not discount the fact that detection remains far from exhaustive for plants and thus should not be overlooked.

### 2. Effects of the ecological variables

Our results showed that species conspicuousness positively affects detection probability, while habitat closure negatively affects it. This general result was expected, as it has also been observed for plants in occupancy and time-to-detection studies (Alexander et al., 2012; Garrard et al., 2008; Hauser et al., 2022). In open habitats, detection probability was the same regardless of the species’ conspicuousness. In contrast, differences between species appeared with increasing habitat closure, reaching up to a 0.42 difference in detection probability between the extreme levels of species conspicuousness in the most closed habitats (estimated difference for the unlimited count with 3 min counting time). It shows that habitat closure drives the detection probability of plant individuals more than species conspicuousness. Furthermore, since both species conspicuousness and habitat closure usually vary over time and space, our results show that heterogeneous detection probability is common and should be considered as the rule rather than the exception. Estimating temporal or spatial trends using raw counts when detection is heterogeneous is likely to yield biased estimates and should therefore be avoided by all means.

The negative effect of the density of individuals on detection probability was unexpected in its magnitude (e.g. a predicted detection probability of 0.67 in a quadrat with 20 individuals to 0.37 in a quadrat with 300 individuals, using the unlimited count in average conditions) and occurred regardless of the counting method. Although it would have supported our results, we did not find this phenomenon reported in the literature, even for animal taxa, where density-dependent detection probabilities usually result from changes in individual behaviour in response to density variations (Veech et al., 2016), and not to the observer detecting fewer individuals. In our case, it may be due to the Weber–Fechner law regarding how humans perceive numbers (see Nieder, 2020, for a summary). Indeed, studies on human cognition have shown that it becomes increasingly difficult to estimate the number of items in an area as the number grows, and that there is a general tendency to underestimate it. It is plausible that this phenomenon seldom applies to animal counts, as these do not usually involve counting immobile items in high density, as is the case for plants.

### 3. Observer effect

The experience in botany of observers had little effect on detection probability compared to the ecological variables and the counting time. In average conditions, the difference in detection probability between novices and the most experienced observers was 0.04 for the quick count, 0.09 for the unlimited count, and 0.17 for the cell count, and confidence intervals largely overlapped except for the cell count (Appendix 6). The advantages of having experience in botany when searching for individuals of a target species may be the ability to more quickly identify an individual once it has been detected, to make fewer identification errors, and possibly to survey the study area more efficiently by knowing where to search for the species (Archaux et al., 2006). These advantages were probably minor in our experiment, since observers were trained to identify the target species before starting counts, all species were easy to identify, and an efficient surveying strategy was of little use in 1×1 m quadrats. In this context, the observers’ task can be seen as searching for small objects in a noisy environment, and it seems plausible that their performance was mainly driven by their ability to remain focused during a repetitive task, their motivation, or their fatigue during the experiment. Larger differences between observers might have been observed in larger quadrats, as the surveying strategy might then be important. Overall, it is likely that the variables we studied have a similar effect on the detection probability regardless of quadrat size. In larger quadrats, the observation process has an additional layer of complexity as observers have to walk in the quadrat and might follow different surveying strategies. Furthermore, the differences between observers might have been greater if we had involved participants with recent experience in searching for the target species (Hauser et al., 2022) due to the well-known ‘search image’ effect, whereby foraging predators improve their probability of detecting the type of prey they are trained to detect (Ishii & Shimada, 2009). If this effect indeed applies to plant counts, population trends estimated from raw counts will be biased if the same observers always make the counts, as their detection probability will increase with time.

### 4. Comparison of the counting methods

As expected, increasing the survey effort, measured by the time spent counting, massively increased detection probability (Archaux et al., 2006; Hauser et al., 2022). In average conditions, the estimated detection probability was 0.36 for the quick count (30 sec counting time), 0.54 for the unlimited count (1.8 min average counting time), and 0.73 for the cell count (8.2 min). In addition, for the unlimited and cell counts, observers spending more time counting achieved a higher detection probability (e.g. for the unlimited count in average conditions, detection probability increased from 0.50 at 1 min to 0.83 at 10 min).

The gain in detection associated with counting individuals per cell instead of the whole quadrat could be due to two reasons: the cells force the observer to survey the whole surface area of the quadrat uniformly, and might also reduce uncertainty whether or not an individual has already been counted, which frequently occurs when individuals are close to each other. However, contrary to our expectations, increasing survey effort did not reduce the effect of the ecological variables on detection probability, the most striking being the density of individuals, which had an almost identical effect on detection probability for all three counting methods.

### 5. Unexplained variance and implications of detection probability heterogeneity

Of the random effects included in the model, the most unexplained variation in detection probability was at the inter-session and inter-quadrat levels. Since we only measured three synthetic variables to characterise the differences between sessions and quadrats (species conspicuousness, habitat closure and true density), we expected that unexplained variance would remain at these levels. For example, we did not measure colour and height heterogeneity, for the habitat or for the target species, although these variables certainly impact detection probability (Garrard et al., 2013; Hauser et al., 2022).

The fact that the unexplained inter-observer variation was the lowest of the random effects is an encouraging result for future studies, as it means that there are few differences in detection probability between observers compared to the differences between species or over a habitat closure gradient. Yet, overall, more than half (65%) of the variance in detection probability remained unexplained by our model. Thus, observers had a heterogeneous detection probability between quadrats even after controlling for several variables, suggesting that factors that vary over time for each observer, such as concentration or motivation, probably explain a large part of the variation in detection probability.

### 6. Recommendations for counting plants

Our study shows for plants what has been documented many times for animals: that detection probability of individuals is less than one and is usually heterogeneous in space and time, regardless of the counting method and survey effort. Using counts affected by imperfect detection as a proxy for the density of individuals to make comparisons over time and space is only valid under the hypothesis of constant detection. Our findings show that multiple ecological and observational variables cause heterogeneity in detection probability. While observational variables can be standardised, it seems unlikely that ecological variables (i.e. species conspicuousness, habitat closure and true density of individuals) will be constant in any study, even those conducted at limited temporal and spatial scales. Furthermore, trying to achieve almost perfect detection (i.e. >0.95) by increasing survey efforts to fix the issue of detection heterogeneity seems unrealistic given the resource constraints of most ecology and conservation studies. As we studied a wide variety of species and habitats, we trust that our results are transferable to most biomes and species types, provided the scientific context is comparable. The spatial scale must be equivalent in terms of quadrat size (i.e. small enough that it is not necessary to walk inside during the survey) and the studied species must be of a similar life form (e.g. herbaceous species or tree seedlings). We therefore argue that temporal or spatial trends in plant populations should not be estimated using raw counts, as they could then reflect a trend in ecological variables rather than the population’s fate.

The detection probability depends on the conditions of the observation process and the studied system; thus, it has to be estimated within each study and cannot be extrapolated from our results. Several methods can be used to estimate individual detection probability, and thus estimate unbiased population sizes or trends (e.g. Lebreton et al., 1992; Buckland et al., 2001). In particular, the N-mixture method uses repeated counts of unmarked individuals on sampling units (e.g. quadrats or sites) to estimate detection probability from the differences in the number of individuals detected between counts (Royle, 2004). With this method, estimates of population sizes or trends are more precise if detection probability is high and homogeneous in time and space, if the mean density of individuals is high, and if the sample size, i.e. the number of sampling units and count repetitions, is high (Royle, 2004; Ficetola et al., 2018). Thus, for studies using the N-mixture method, no matter the studied population and the counting method used, we recommend setting a fixed counting time as leaving it to the observer’s choice will add heterogeneity to detection probability, resulting in less precise estimates.

In terms of allocating survey effort, there are three options: use a more time-consuming counting method to increase detection probability, perform more count repetitions, or survey more sampling units. The only way to be sure of the optimal design is to conduct a pilot study to estimate the mean density of individuals and their detection probability in the population of interest. However, Ficetola et al. (2018) have shown that when the detection probability is above 0.10, the most advantageous strategy is usually to make three count repetitions and to increase the number of sampling units as much as possible. Conversely, when the detection probability is below 0.10, it is better to increase it by using a more time-consuming counting method or to increase the number of count repetitions. Thus, we recommend counting individuals in quadrats without cells with a fixed short counting time (e.g. 0.5–1 min), as this method will yield a detection probability above 0.10 for most species and allow the most sampling units to be surveyed. More time-consuming counting methods should be used only if the study population grows in a very closed habitat and the individuals are inconspicuous, or if travel costs resulting from increasing the number of sampling units is a strong constraint.

## Acknowledgements

We would like to warmly thank the 167 volunteers (Appendix 9) who participated in the experiment, including during the preliminary tests. We are grateful to the volunteers who helped organise the field sessions by finding study sites or participants, Virginie Pons for her help in transcribing the data, and Solal Boutoux for his help in designing Fig. 1.

## Authors’ contributions

J.P., A.B., A.C. and G.P. conceived the study and designed the methodology; J.P. and G.P. organised and conducted the field sessions; J.P. analysed the data and led the writing of the manuscript. All authors critically contributed to the drafts and gave their final approval for publication.

## Data availability statement

All data and code used for this work are available from the GitHub repository: https://github.com/JanPerret/lndividual_detection_in_plant_counts

