## Appendix for "Plants stand still but hide: imperfect and heterogeneous detection is the rule when counting plants"

Created on the 2022-12-21

### Appendix 1: Summary of the field sessions

**Table S1.1:** Summary of the field sessions.

| **Species** | **Date** | **Locality** | **Predominant phenological stage** | **Habitat** | **Site number** | **Number of participants** |
| --- | --- | --- | --- | --- | --- | --- |
| Plantago lanceolata | 07/10/2020 | Montpellier (34090) | Vegetative | Urban wasteland | 1 | 6 |
| Allium sp. | 16/10/2020 | Montpellier (34090) | Vegetative | Grassland | 2 | 3 |
| Calendula arvensis | 20/10/2020 | Montpellier (34090) | Flowering | Grassland | 2 | 5 |
| Sanguisorba minor | 28/10/2020 | Montpellier (34090) | Vegetative | Urban wasteland | 1 | 5 |
| Allium chamaemoly | 05/02/2021 | Lattes (34970) | Vegetative | Western Mediterranean scrubland | 3 | 6 |
| Euphorbia helioscopia | 04/03/2021 | Saint-Jean-de-Vedas (34430) | Flowering | Dry calcareous grassland | 4 | 4 |
| Erodium cicutarium | 18/03/2021 | Clapiers (34077) | Flowering | Western Mediterranean scrubland | 5 | 6 |
| Chrysosplenium alternifolium | 13/04/2021 | Valjouffrey (38740) | Flowering | Alpine riparian forest | 6 | 5 |
| Fagus sylvatica | 14/04/2021 | Gap (05000) | Seedling | Beech forest | 7 | 6 |
| Hieracium sp. | 15/04/2021 | Le Muy (83490) | Vegetative | Western Mediterranean scrubland | 8 | 7 |
| Bromus rubens | 16/04/2021 | Marseille (13000) | Flowering | Mediterranean grassland | 9 | 6 |
| Sherardia arvensis | 23/04/2021 | Clapiers (34077) | Flowering | Western Mediterranean scrubland | 5 | 6 |
| Mentha suaveolens | 05/05/2021 | Montferrier-sur-Lez (34980) | Vegetative | Lowland hay meadow | 10 | 5 |
| Scabiosa atropurpurea | 06/05/2021 | Clapiers (34077) | Vegetative | Mediterranean pine forest | 11 | 6 |
| Ornitogalum angustifolium | 11/05/2021 | Mas-de-Londres (34380) | Flowering | Mediterranean extensive pasture | 12 | 5 |
| Evax pygmaea | 18/05/2021 | Arles (13200) | Flowering | Mediterranean sand annual grassland | 13 | 7 |
| Ophrys lutea | 19/05/2021 | Mas-de-Londres (34380) | Flowering | Mediterranean extensive pasture | 12 | 5 |
| Carex flacca | 26/05/2021 | Mas-de-Londres (34380) | Vegetative | Mediterranean extensive pasture | 12 | 6 |
| Anacamptis pyramidalis | 27/05/2021 | Mas-de-Londres (34380) | Flowering | Mediterranean extensive pasture | 12 | 5 |
| Rumex acetosella | 07/06/2021 | Orsay (91400) | Flowering | Lowland hay meadow | 14 | 5 |
| Equisetum telmateia | 11/06/2021 | Orsay (91400) | Vegetative | Megaphorb | 15 | 7 |
| Linaria vulgaris | 15/06/2021 | Haulme (08800) | Vegetative | Lowland hay meadow | 16 | 7 |
| Melampyrum arvense | 16/06/2021 | Rancennes (08600) | Flowering | Lowland hay meadow | 17 | 8 |
| Veronica chamaedrys | 17/06/2021 | Haulme (08800) | Flowering | Frequently mown meadow | 18 | 5 |
| Limonium narbonense | 25/06/2021 | Le Sambuc (13200) | Vegetative | Salt meadow | 19 | 5 |
| Erigeron annuus | 28/06/2021 | Creys-Mepieu (38510) | Flowering | Lowland hay meadow | 20 | 5 |
| Ambrosia artemisiifolia | 07/07/2021 | Avignon (84140) | Vegetative | River gravel bank | 21 | 6 |
| Limonium girardianum | 28/07/2021 | Le Grau-du-Roi (30240) | Vegetative | Salt steppe | 22 | 6 |
| Euphorbia peplis | 29/07/2021 | Vic-la-Gardiole (34110) | Flowering | Sand beach | 23 | 7 |
| Epipogium aphyllum | 06/08/2021 | Crots (05200) | Flowering | Pine forest | 24 | 6 |

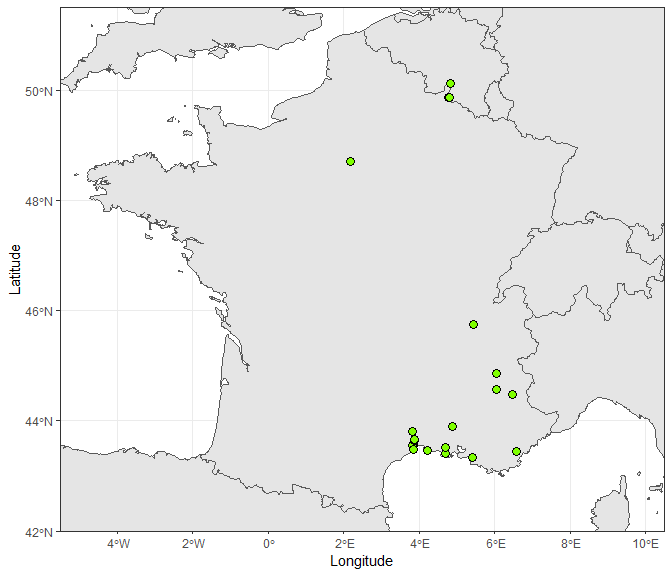

**Figure S1.1:** Map of the 24 study sites. Some sites were only a few kilometres apart; in this case, the corresponding points overlap.

### Appendix 2: Detailed Mat&Meth

#### Creation of the species conspicuousness variable

For each field session, we visually estimated the mean height range of the individuals of the target species and recorded whether or not the majority of individuals were flowering. Inconspicuous flowers (i.e. without petals or without vivid colours) were not taken into account. We then created the species conspicuousness variable with the following criteria:

1 -> Individuals [0 cm; 5 cm] tall, without colourful flowers

2 -> Individuals [5 cm; 10 cm] tall, or [0 cm; 5 cm] with colourful flowers

3 -> Individuals [10 cm; 20 cm] tall, without colourful flowers

4 -> Individuals [20 cm: +Inf[ tall, or [10 cm; 20 cm] with colourful flowers

**Table S2.1:** Value of the species conspicuousness variable for the 30 sampled species (1 = least conspicuous; 4 = most conspicuous).

| **Species name** | **Species code** | **Species conspicuousness** | **Date** |
| --- | --- | --- | --- |
| Allium chamaemoly | ALLCHA | 1 | 05/02/2021 |
| Bromus rubens | BRORUB | 1 | 16/04/2021 |
| Carex flacca | CARFLA | 1 | 26/05/2021 |
| Erodium cicutarium | EROCIC | 1 | 18/03/2021 |
| Euphorbia peplis | EUPPEP2 | 1 | 29/07/2021 |
| Evax pygmaea | EVAPYG | 1 | 18/05/2021 |
| Fagus sylvatica | FAGSYL | 1 | 14/04/2021 |
| Limonium girardianum | LIMGIR | 1 | 28/07/2021 |
| Plantago lanceolata | PLALAN | 1 | 07/10/2020 |
| Sherardia arvensis | SHEARV | 1 | 23/04/2021 |
| Allium sp | ALLsp | 2 | 16/10/2020 |
| Ambrosia artemisiifolia | AMBART | 2 | 07/07/2021 |
| Calendula arvensis | CALARV2 | 2 | 20/10/2020 |
| Chrysosplenium alternifolium | CHRALT | 2 | 13/04/2021 |
| Epipogium aphyllum | EPIAPH | 2 | 06/08/2021 |
| Hieracium sp | HIEsp | 2 | 15/04/2021 |
| Ophrys lutea | OPHLUT | 2 | 19/05/2021 |
| Rumex acetosella | RUMACE | 2 | 07/06/2021 |
| Sanguisorba minor | SANMIN | 2 | 28/10/2020 |
| Scabiosa atropurpurea | SCAATR | 2 | 06/05/2021 |
| Euphorbia helioscopia | EUPHEL | 3 | 04/03/2021 |
| Limonium narbonense | LIMNAR | 3 | 25/06/2021 |
| Linaria vulgaris | LINVUL | 3 | 15/06/2021 |
| Melampyrum arvense | MELARV | 3 | 16/06/2021 |
| Ornitogalum angustifolium | ORNANG | 3 | 11/05/2021 |
| Veronica chamaedrys | VERCHA | 3 | 17/06/2021 |
| Anacamptis pyramidalis | ANAPYR | 4 | 27/05/2021 |
| Equisetum telmateia | EQUTEL | 4 | 11/06/2021 |
| Erigeron annuus | ERIANN | 4 | 28/06/2021 |
| Mentha suaveolens | MENSUA | 4 | 05/05/2021 |

#### Creation of the habitat closure variable

For each quadrat, we visually estimated the proportion of the area covered by vegetation, using the cells as a guide. We also measured the height of the vegetation at 9 identical points in each quadrat, marked by the intersections of the elastic bands forming the cells (Fig. S2.1). The median of these 9 measurements was used as a measure of the height of the vegetation in each quadrat.

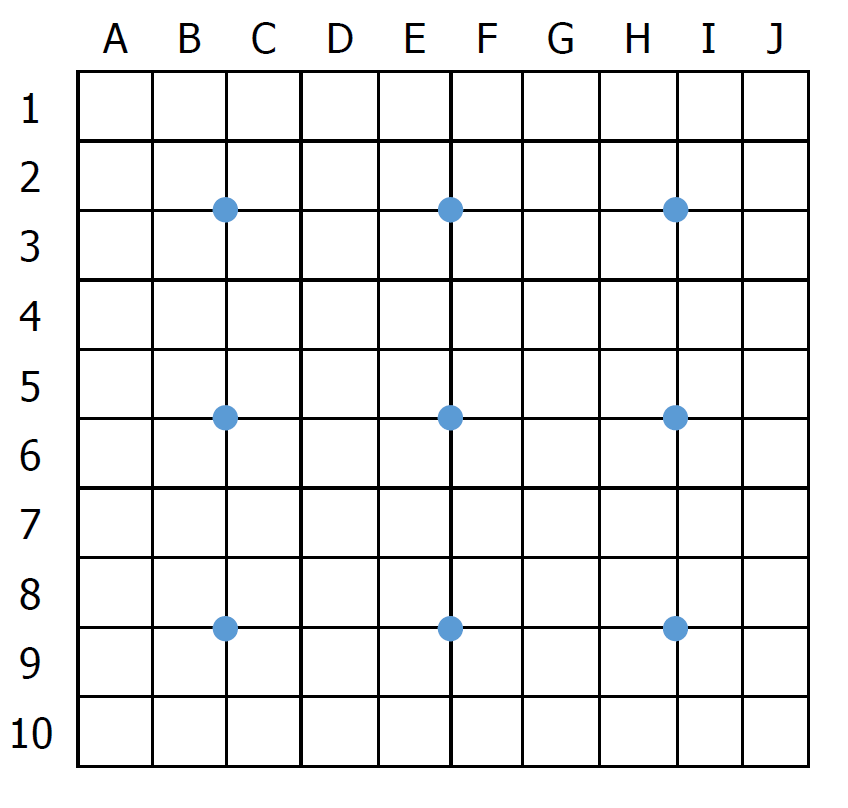

**Figure S2.1:** Diagram with the location of the points (in blue) where we measured vegetation height in each quadrat.

In order to obtain a synthetic measure of vegetation closure, we performed a PCA on the two variables (i.e. vegetation cover proportion and vegetation height) and kept the quadrat coordinates on the first principal component.

The first principal component explained 77.18% of the total variance, and the second explained 22.82% of the variance. Figure S2.2 shows the value associated with each quadrat on the first principal component, i.e. our new composite variable of habitat closure, as a function of the cover and height of the vegetation.

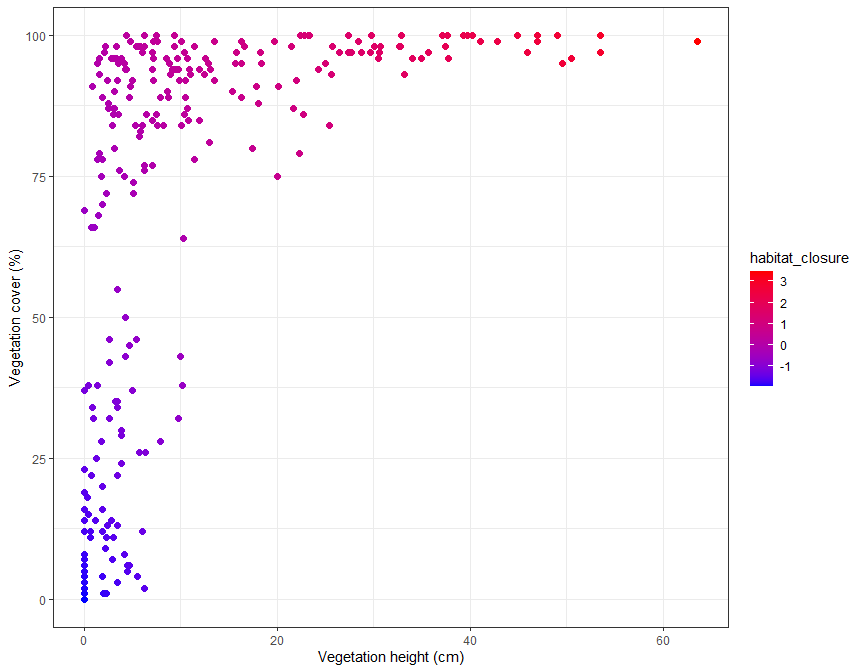

**Figure S2.2:** Value of the ‘habitat closure’ variable as a function of vegetation height and cover (-1 = least closed; 3 = most closed).

#### Evaluation of the experience in botany of the observers

Before we explained the experiment to the observers, they were asked to fill in a form including a rating of their level of experience in botany on a scale of 1 to 10. In order to standardise the ratings between observers of all field sessions, a brief description was provided for every second level of this experience scale. We subsequently divided the experience level by two to obtain a scale with 5 classes (1 = least experienced; 5 = most experienced).

The form was as follows:

**How do you evaluate your experience in botany? (Circle your level)**

The number of known species is given as a guide. If the description of a level fits, but you can easily recognise more species than the indicated range, you can choose the higher category if you feel that it better fits your experience level. A ‘known species’ means that you can identify it with certainty without using an identification guide, or simply by quickly checking a single criterion.

1

2: You have previously practised botany but rarely do so now or have not practised at all for a long time. (Less than 50 known species)

3

4: You practice botany occasionally, or you started less than 3 years ago. (50–300 known species)

5

6: You practice botany regularly and have done so for more than 3 years. (300–1000 known species)

7

8: You have been practising botany intensively for more than 5 years and are developing expertise. (1000–2000 known species)

9

10: You have been practising botany intensively for more than 10 years and are recognised as an expert. (More than 2000 known species)

### Appendix 3: Data visualisation

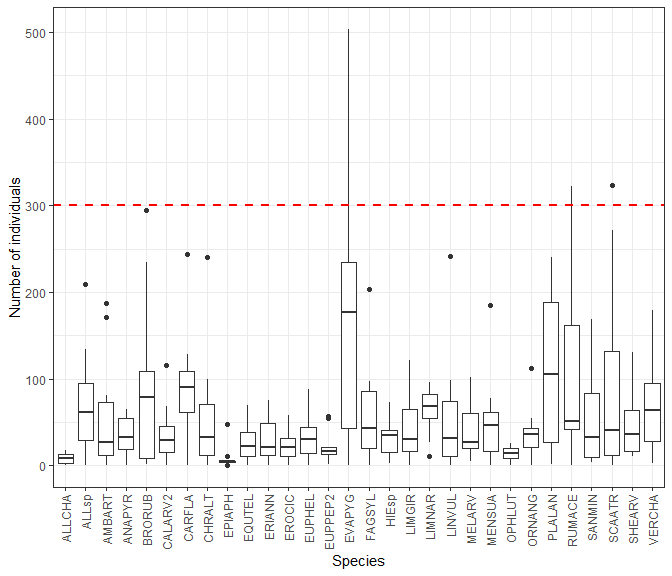

**Figure S3.1:** The true number of individuals per quadrat and field session, before the four density outliers were removed. The dotted red line shows the threshold of 300 individuals above which we removed the quadrats from the dataset to avoid the leverage effect.

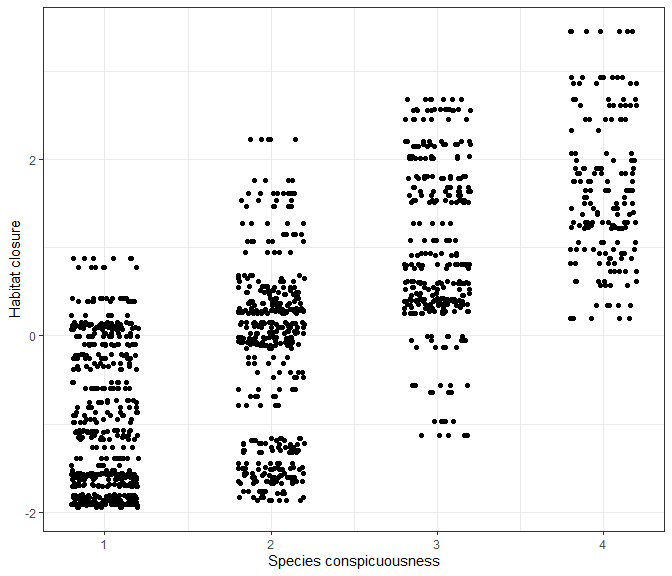

**Figure S3.2:** Distribution of the observations depending on the conspicuousness of the studied species and the habitat closure. Each point represents an observation by a participant on a quadrat for the first counting method, i.e. only one-third of the observations are represented. Random noise was added to the position of the points on the x-axis so that they do not overlap. The combinations that do not exist in the dataset, i.e. a very inconspicuous species in a very closed habitat or a very conspicuous species in a very open habitat, are uncommon in the field.

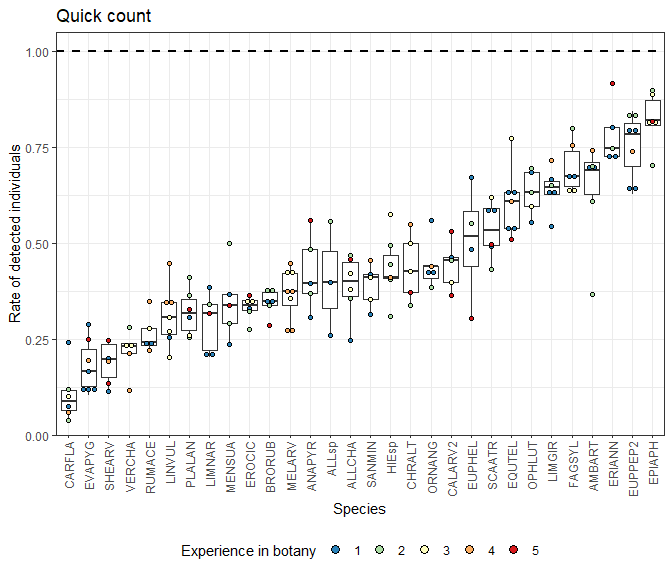

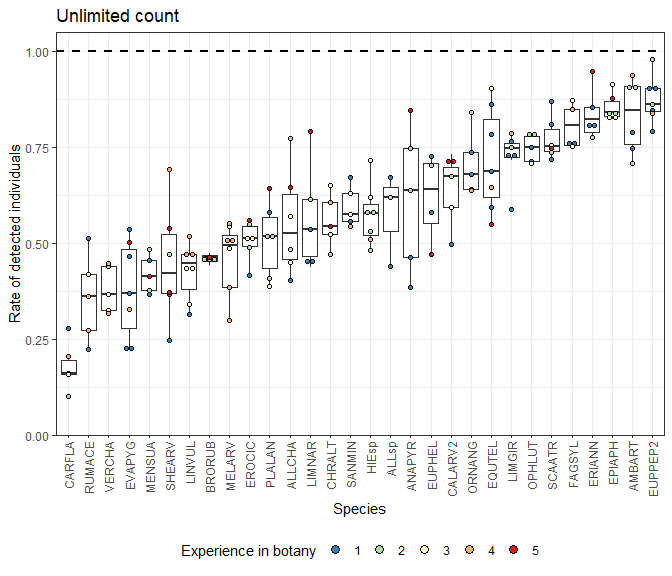

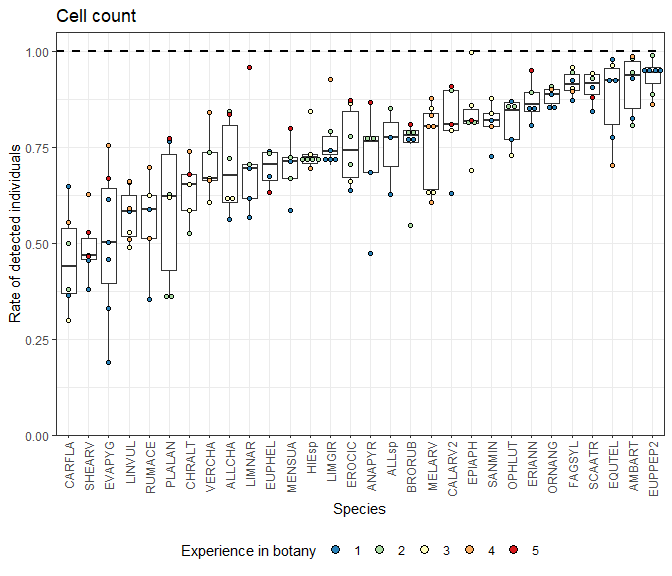

**Figure S3.3:** Mean proportion of detected individuals per observer after the excess detections were removed. Each point represents the mean detection proportion of an observer during a field session, with 1.00 being perfect detection. The colours indicate the observer’s experience in botany (1 = least experienced; 5 = most experienced).

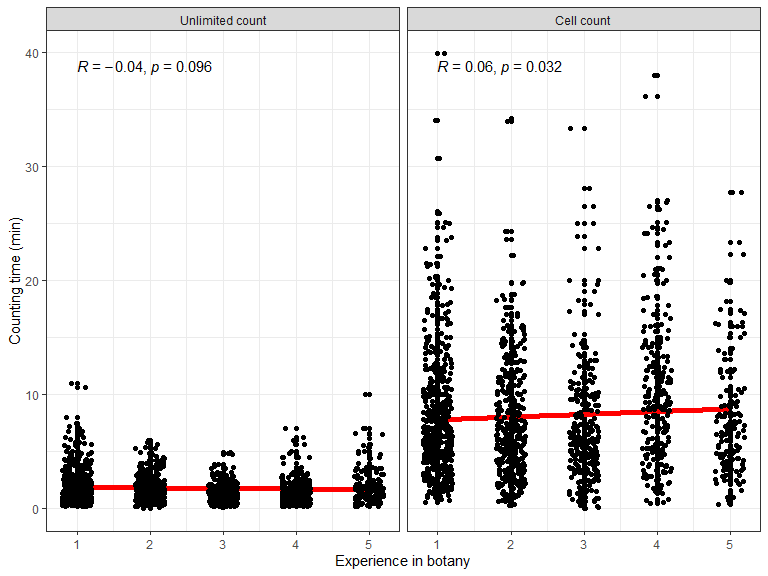

**Figure S3.4:** Distribution of the counting times depending on the level of experience in botany of the observers. Random noise was added to the position of the points on the x-axis so that they do not overlap. The red lines show basic linear regressions, and the values written in the top left corner are Pearson’s correlation coefficients and the p-value of the associated test.

### Appendix 4: Description of the excess detections

The dataset was composed of a total of 5024 observations before removing the 34 quadrats that did not contain any individual of the target species. 565 observations were made in these empty quadrats, only 12 of which with excess detections. This seems to indicate that identification errors were extremely rare during the experiment, and that the excess detections were mainly due to errors in differentiating individuals.

The description below refers to the dataset after removing the quadrats that did not contain any individual of the target species. After this filtering, the dataset contained 4459 observations, of which 140 excess detections.

**Table S4.1:** Distribution of the observations with excess detections per field session.

| **Species** | **Total number of observations** | **Number of excess detections** | **Proportion of excess detections** |
| --- | --- | --- | --- |
| ALLCHA | 144 | 2.000 | 0.014 |
| ALLsp | 80 | 0.000 | 0.000 |
| AMBART | 144 | 6.000 | 0.042 |
| ANAPYR | 120 | 1.000 | 0.008 |
| BRORUB | 180 | 6.000 | 0.033 |
| CALARV2 | 135 | 4.000 | 0.030 |
| CARFLA | 143 | 0.000 | 0.000 |
| CHRALT | 120 | 7.000 | 0.058 |
| EPIAPH | 144 | 0.000 | 0.000 |
| EQUTEL | 180 | 7.000 | 0.039 |
| ERIANN | 120 | 32.000 | 0.267 |
| EROCIC | 162 | 1.000 | 0.006 |
| EUPHEL | 108 | 6.000 | 0.056 |
| EUPPEP2 | 210 | 9.000 | 0.043 |
| EVAPYG | 122 | 0.000 | 0.000 |
| FAGSYL | 161 | 11.000 | 0.068 |
| HIEsp | 210 | 10.000 | 0.048 |
| LIMGIR | 162 | 6.000 | 0.037 |
| LIMNAR | 146 | 2.000 | 0.014 |
| LINVUL | 168 | 1.000 | 0.006 |
| MELARV | 240 | 1.000 | 0.004 |
| MENSUA | 120 | 0.000 | 0.000 |
| OPHLUT | 120 | 7.000 | 0.058 |
| ORNANG | 135 | 0.000 | 0.000 |
| PLALAN | 180 | 8.000 | 0.044 |
| RUMACE | 105 | 0.000 | 0.000 |
| SANMIN | 150 | 2.000 | 0.013 |
| SCAATR | 126 | 8.000 | 0.063 |
| SHEARV | 175 | 2.000 | 0.011 |
| VERCHA | 149 | 1.000 | 0.007 |

The observations with excess detections made up less than 6.8% of the observations per field session, except for the session on Erigeron annuus (ERIANN; 26.7% observations over 100% detection). For this field session the detection of individuals was very easy as the individuals were in full flower and taller than the surrounding vegetation, which resulted in a high detection rate. Furthermore, many individuals had multiple stems and the ramification was close to the ground. It was possible to differentiate the individuals unambiguously by looking closely at the ramification, but the observers did not pay close attention. This combination of easy detection and the particular morphology of individuals led to these excess detections.

**Table S4.2:** Distribution of the observations with excess detections per level of botany experience (1 = least experienced; 5 = most experienced).

| **Experience in botany** | **Total number of observations** | **Number of excess detections** | **Proportion of excess detections** |
| --- | --- | --- | --- |
| 1.000 | 1,474 | 57.000 | 0.039 |
| 2.000 | 1,095 | 28.000 | 0.026 |
| 3.000 | 790 | 25.000 | 0.032 |
| 4.000 | 641 | 14.000 | 0.022 |
| 5.000 | 459 | 16.000 | 0.035 |

The excess detections are roughly evenly distributed across the levels of experience in botany.

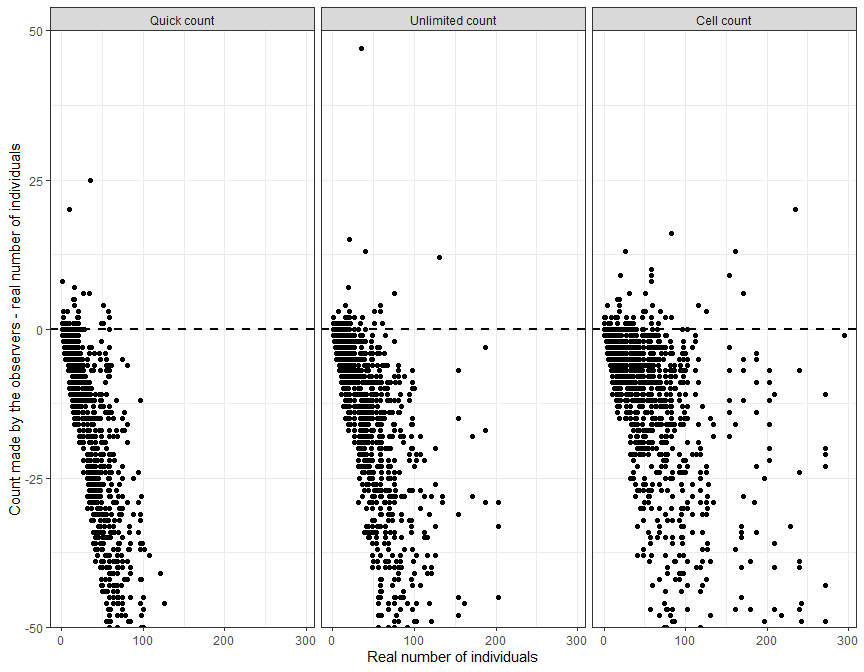

**Figure S4.1:** Difference between the counts made by the observers and the true number of individuals present in the quadrats. The lower part of the graph is truncated, but not the upper part, so all excess detections are represented. The majority of the excess detections occurred in quadrats containing few individuals, and were below 110% of the true number of individuals.

### Appendix 5: Model goodness-of-fit assessment

We assessed the goodness-of-fit of the model using the R package DHARMa (Hartig, 2022). This package uses a simulation approach to produce scaled residuals for Generalised Linear Mixed Models (GLMMs), which can be interpreted as residuals from classic linear models to assess a model’s fit. For a detailed explanation of how the package works and how to interpret residual diagnostic plots, see the vignette of the package here: <https://cran.r-project.org/web/packages/DHARMa/vignettes/DHARMa.html>

#### Main residual diagnostic plots

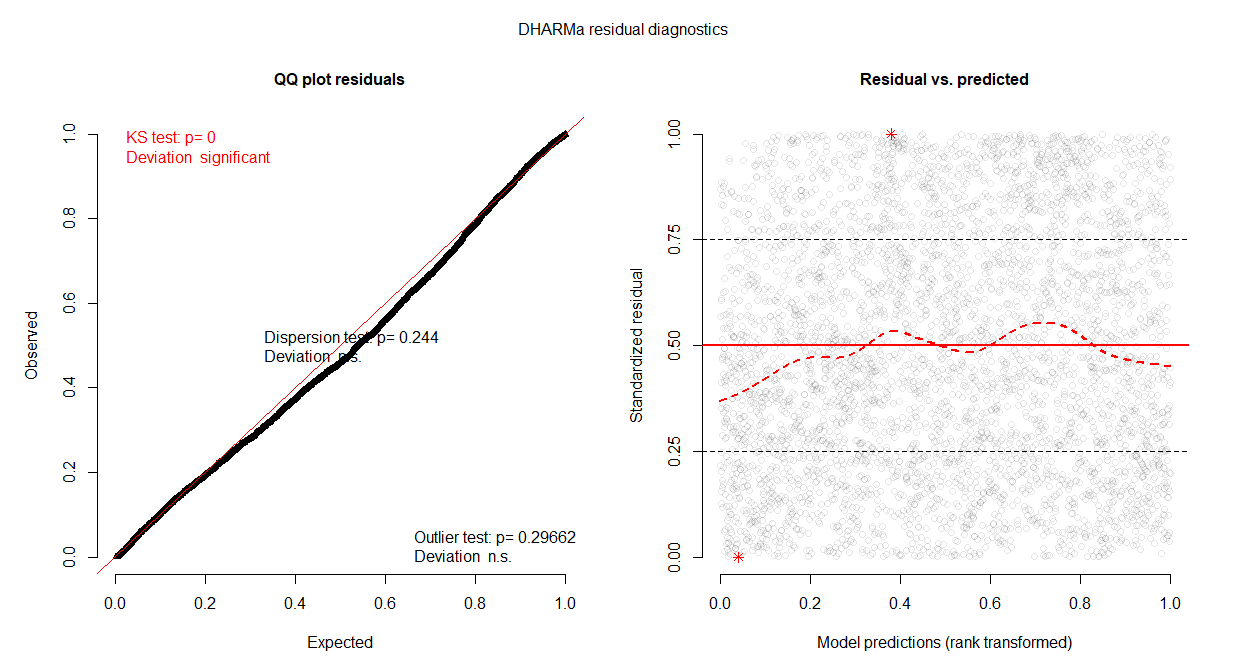

Overall the model fit seems very good! On the quantile-quantile plot (left) there is a slight deviation from the x=y line, but this deviation is very small. On the three tests performed by the DHARMa package, the dispersion test and the outlier test are non-significant, which means that the simulated dispersion is equal to the observed dispersion and that there are no simulation outliers, respectively. However the KS uniformity test is significant, which seems logical given the deviation we can see on the QQ plot and the large number of observations in the dataset. A visual inspection of the QQ plot shows that although the deviation exists, it is small. It seems unlikely that this could substantially affect the model estimates, especially given the robustness of GLMMs to violations of distributional assumptions (Schielzeth et al., 2020). The residuals against predicted values plot (right) supports this interpretation, as there is no clear pattern in the residuals.

#### Residuals against predictors plots

We now plot the residuals against the predictors in the model to check if there is a lack of fit for some predictor.

##### Species conspicuousness

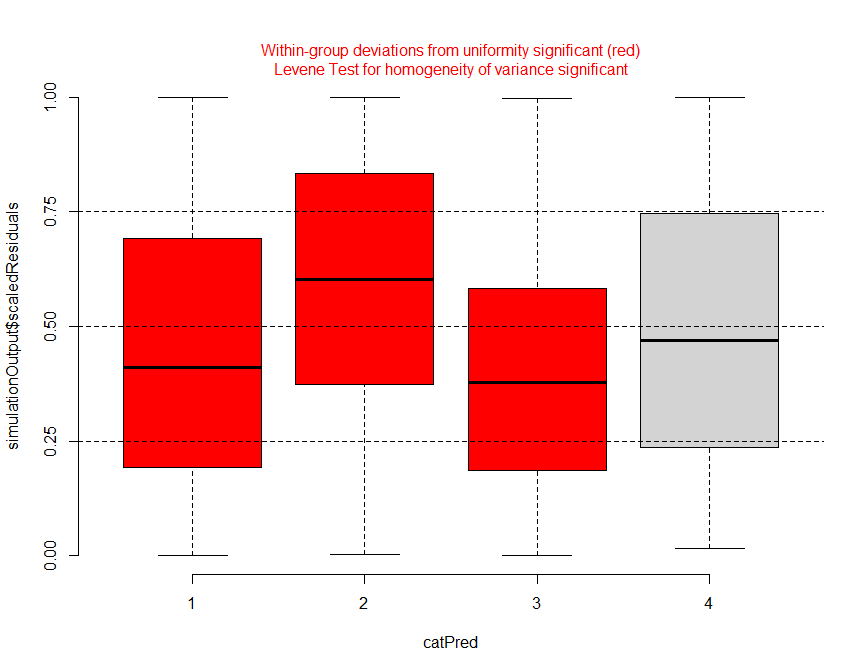

There is a slight deviation from the expected distribution, but no systematic pattern.

##### Habitat closure

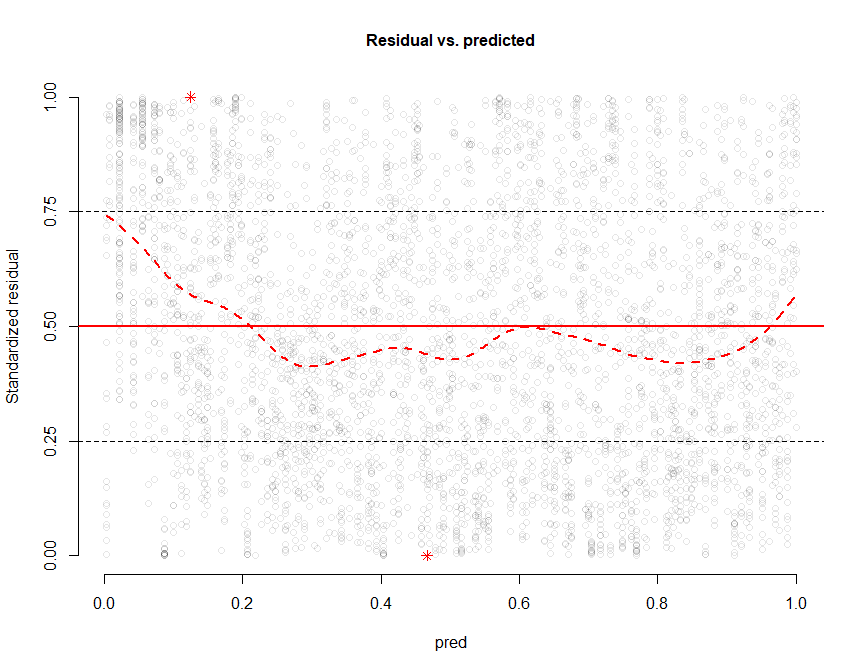

There is a slight deviation from a perfect uniform distribution as it can be seen by the fitted quantile regression line (dashed red line), but it seems small and without a systematic pattern.

##### Counting method

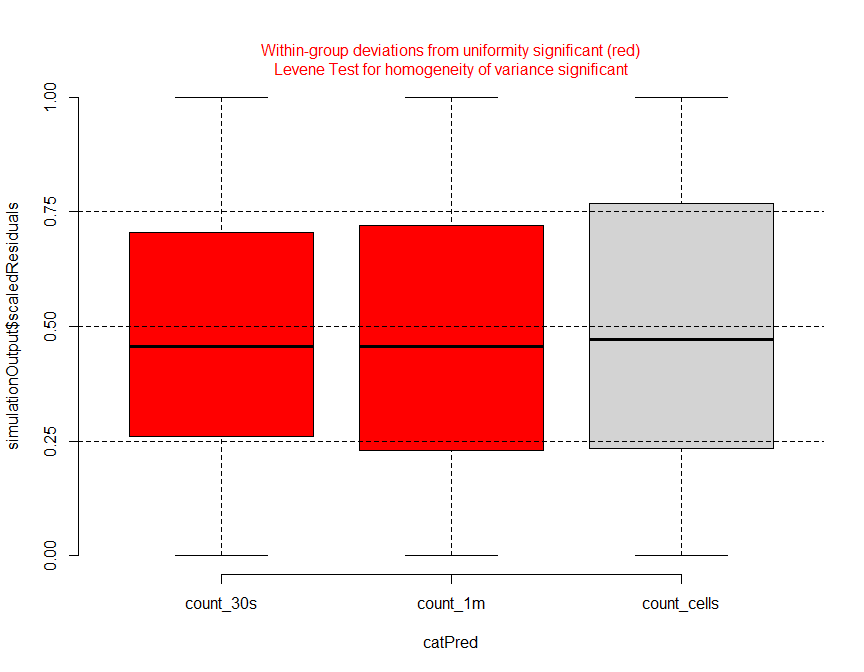

There is a statistically significant deviation from a uniform distribution for the quick count and the unlimited count (count_30s and count_1m, respectively), but the box-whisker plot shows that this deviation is very small.

##### Experience in botany

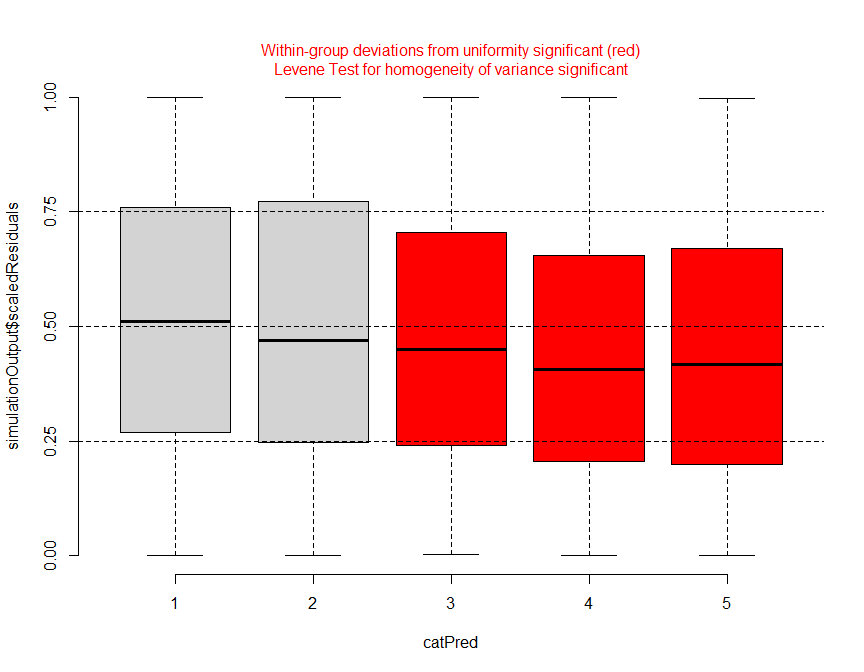

As for the counting methods, there is a deviation for the experience levels in botany 3, 4 and 5. There could be a systematic pattern here (deviation increases with the level in botany), but the deviation is very small, so it seems there is nothing to worry about.

##### True density of individuals

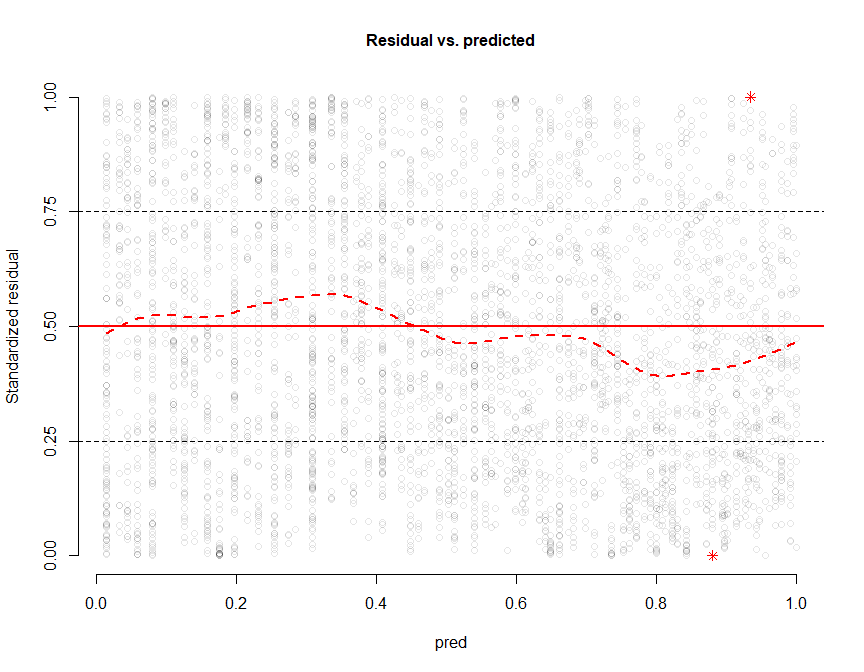

The fitted quantile regression finds a significant deviation but it’s very small and no pattern can be seen in the residuals.

##### Quadrat order

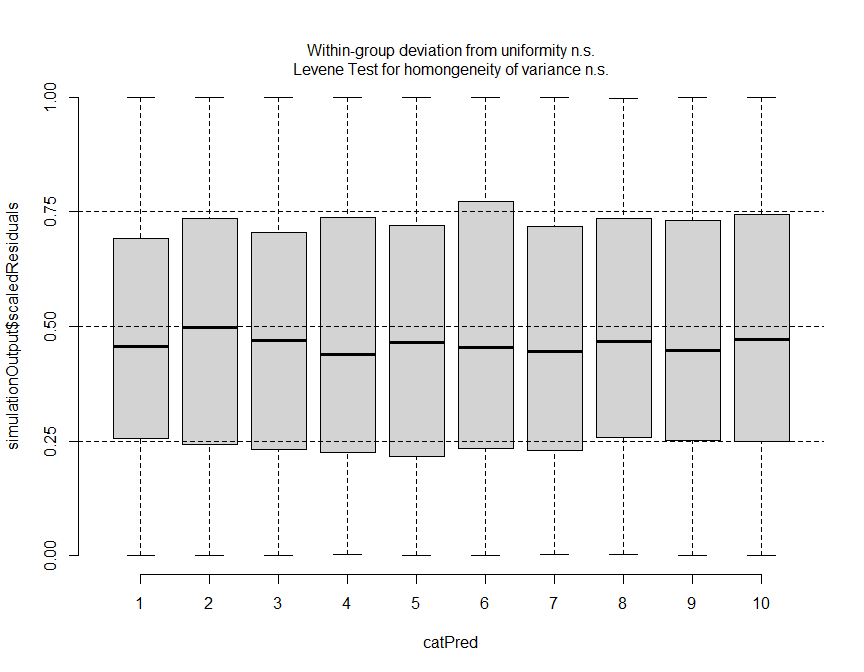

There is no deviation from the expected uniform distribution.

##### Counting time

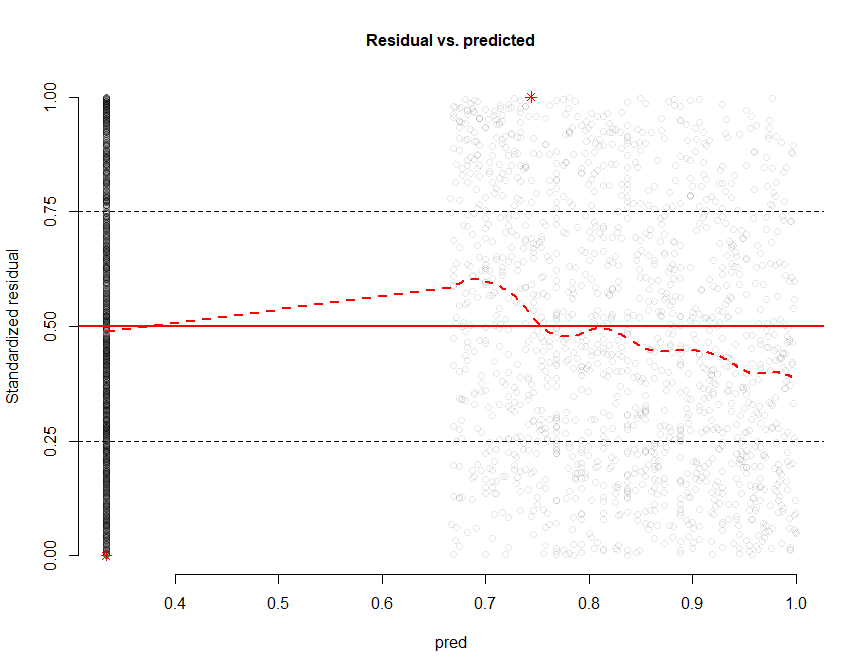

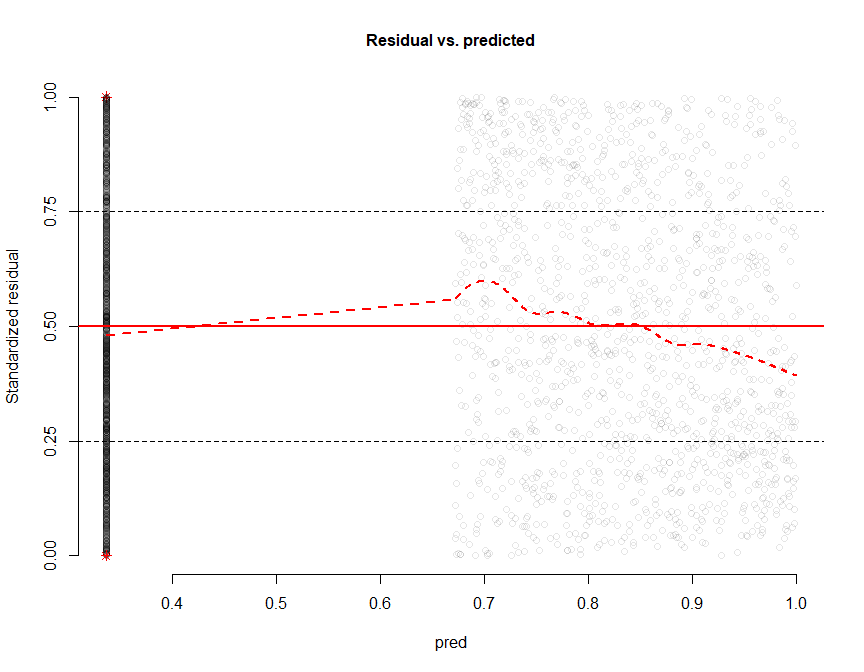

For the two plots above, the strange pattern in the residuals (no points in the left half of the plots) comes from the fact that we had to code manually the two counting time variables, as for one of the three counting methods the counting time was fixed at 30 seconds and not variable as it was for the other two methods. We created two separate variables (counting_time_1m and counting_time_cells), which contained the recorded counting times for the unlimited count and the cell count, respectively. For the rows of the dataset concerning the other counting methods, the values of these two variables were set to zero. Thus, only the right part of the plots have to be interpreted here, and there seems to be no clear pattern in the residuals in this part of the plots.

### Appendix 6: Coefficients of the model

All predictors were standardised before fitting the model. The model fitting function (using the R package lme4) is below:

```
mod_final <- glmer(prop_detect ~ (1|species) + (1|species:quadrat_id) + (1|obs_id)
                   + species_conspicuousness * habitat_closure
                   + species_conspicuousness * counting_method
                   + habitat_closure * counting_method
                   + exp_bota * counting_method
                   + count_TRUE * counting_method
                   + quadrat_order * counting_method
                   + counting_time_1m * habitat_closure
                   + counting_time_1m * species_conspicuousness
                   + counting_time_cells * species_conspicuousness
                   + counting_time_cells * habitat_closure,
                   family = binomial, data = df_3_methods, weight = count_TRUE)
```

**Table S6.1:** Fixed effect coefficients of the selected model.

| **term** | **estimate** | **std.error** | **conf.low** | **conf.high** | **statistic** | **p.value** |
| --- | --- | --- | --- | --- | --- | --- |
| Intercept | -0.0047 | 0.1684 | -0.3347 | 0.3252 | -0.0281 | 0.9776 |
| Species conspicuousness | 0.2103 | 0.1716 | -0.1260 | 0.5467 | 1.2257 | 0.2203 |
| Habitat closure | -0.3102 | 0.1050 | -0.5159 | -0.1045 | -2.9552 | 0.0031 |
| Method [unlimited count] | 0.4256 | 0.0199 | 0.3866 | 0.4646 | 21.3750 | 0.0000 |
| Method [cell count] | 0.8436 | 0.0254 | 0.7938 | 0.8933 | 33.2091 | 0.0000 |
| Experience botany | 0.0595 | 0.0358 | -0.0106 | 0.1295 | 1.6630 | 0.0963 |
| Density | -0.2907 | 0.0488 | -0.3864 | -0.1949 | -5.9519 | 0.0000 |
| Quadrat order | 0.0127 | 0.0082 | -0.0035 | 0.0288 | 1.5381 | 0.1240 |
| Counting time x Method [unlimited count] | 0.2088 | 0.0085 | 0.1921 | 0.2254 | 24.6281 | 0.0000 |
| Counting time x Method [cell count] | 0.4790 | 0.0120 | 0.4555 | 0.5026 | 39.8179 | 0.0000 |
| Species conspicuousness x Habitat closure | 0.1430 | 0.0911 | -0.0356 | 0.3216 | 1.5694 | 0.1166 |
| Species conspicuousness x Method [unlimited count] | 0.0685 | 0.0268 | 0.0159 | 0.1210 | 2.5541 | 0.0106 |
| Species conspicuousness x Method [cell count] | 0.0348 | 0.0325 | -0.0290 | 0.0986 | 1.0702 | 0.2845 |
| Habitat closure x Method [unlimited count] | -0.1246 | 0.0204 | -0.1646 | -0.0845 | -6.1021 | 0.0000 |
| Habitat closure x Method [cell count] | 0.1279 | 0.0230 | 0.0829 | 0.1730 | 5.5639 | 0.0000 |
| Method [unlimited count] x Experience botany | 0.0637 | 0.0112 | 0.0417 | 0.0857 | 5.6836 | 0.0000 |
| Method [cell count] x Experience botany | 0.2072 | 0.0123 | 0.1831 | 0.2313 | 16.8792 | 0.0000 |
| Method [unlimited count] x Density | 0.0237 | 0.0093 | 0.0054 | 0.0420 | 2.5389 | 0.0111 |
| Method [cell count] x Density | 0.0275 | 0.0109 | 0.0062 | 0.0488 | 2.5351 | 0.0112 |
| Method [unlimited count] x Quadrat order | 0.0641 | 0.0114 | 0.0418 | 0.0864 | 5.6414 | 0.0000 |
| Method [cell count] x Quadrat order | 0.0903 | 0.0122 | 0.0665 | 0.1142 | 7.4255 | 0.0000 |
| Habitat closure x Counting time x Method [unlimited count] | -0.0221 | 0.0095 | -0.0407 | -0.0035 | -2.3266 | 0.0200 |
| Species conspicuousness x Counting time x Method [unlimited count] | 0.0068 | 0.0096 | -0.0120 | 0.0257 | 0.7091 | 0.4782 |
| Species conspicuousness x Counting time x Method [cell count] | 0.0513 | 0.0117 | 0.0284 | 0.0742 | 4.3962 | 0.0000 |
| Habitat closure x Counting time x Method [cell count] | -0.2045 | 0.0091 | -0.2222 | -0.1867 | -22.5880 | 0.0000 |

**Table S6.2:** Random effect standard deviations and variances of the selected model.

| **term** | **SD** | **variance** |
| --- | --- | --- |
| species:quadrat_id | 0.6474 | 0.4191 |
| obs id | 0.3989 | 0.1592 |
| species | 0.7650 | 0.5852 |

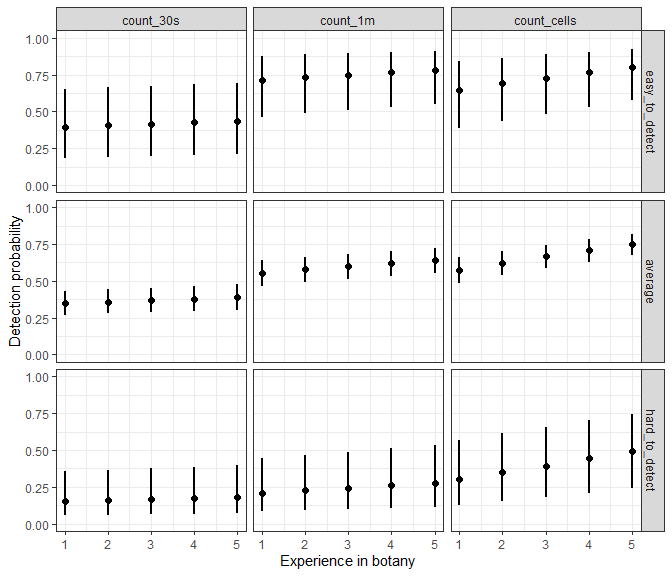

**Figure S6.1:** Predicted detection probability depending on the level of experience in botany of the observer in easy detection conditions (species conspicuousness = 4; habitat closure = -1.5), average conditions (species conspicuousness = 2.5; habitat closure = 0.5) and difficult conditions (species conspicuousness = 1; habitat closure = 2.5). Only the values in average conditions are used in the main text.

### Appendix 7: Results with excess detections reduced to 100% detection

The three figures below are the alternative versions of Figs 2–4 from the main text, made by fitting the model to the dataset in which the excess detections were set to a 100% detection rate instead of removing them from the dataset. The differences are imperceptible, which indicates that the excess detections did not play a role in the analysis results.

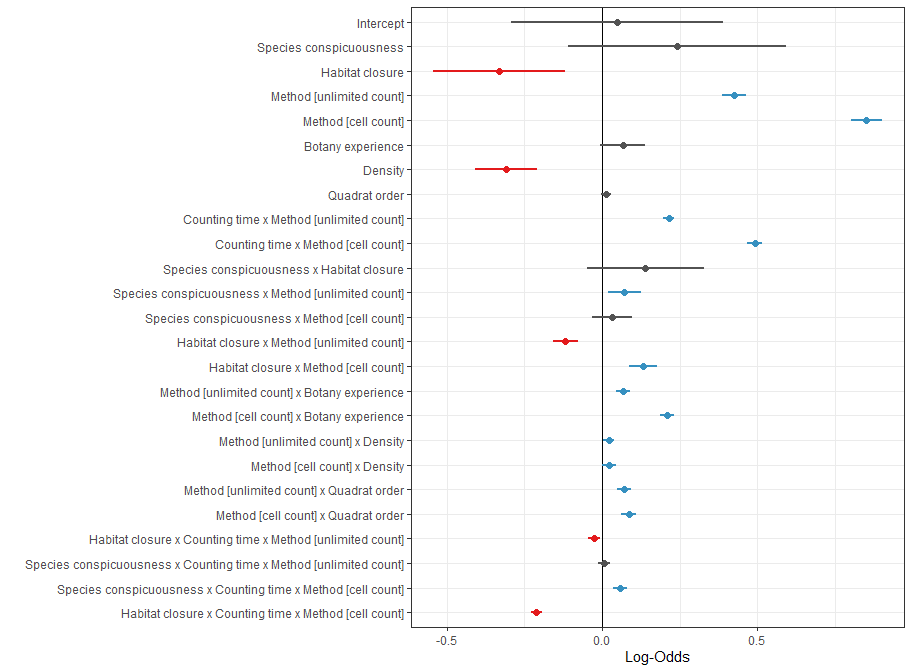

**Figure S7.1:** Coefficients of standardised variables in the selected model. Dots represent estimated coefficients, and lines represent 95% confidence intervals. Positive coefficients (blue) indicate that when the variable increases, the proportion of detected individuals increases, while negative coefficients (red) indicate the opposite. Coefficients with a non-significant effect are shown in grey. For the variable ‘counting method’, the reference level is the estimate of the number of individuals in 30 sec. The coefficients presented for the other two methods indicate the difference in mean detection rate obtained with those methods compared to the reference level.

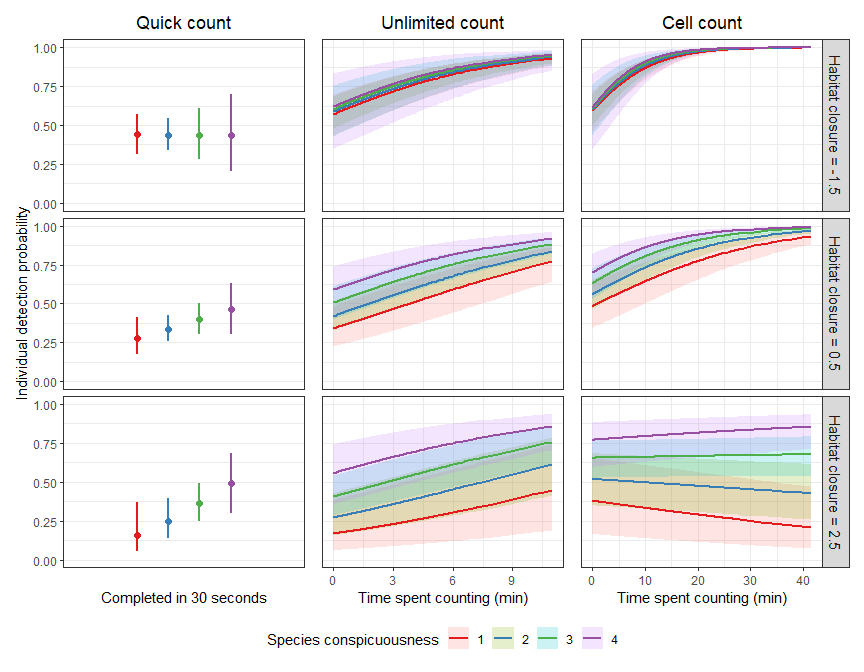

**Figure S7.2:** Predicted detection probability of individuals depending on species conspicuousness, habitat closure, counting time and the counting method used. The other variables included in the model are held constant at the following levels: exp_bota = 2.5, count_TRUE = 100, quadrat_order = 5. Lines represent the mean proportion of detected individuals, and the shading represents 95% confidence intervals.

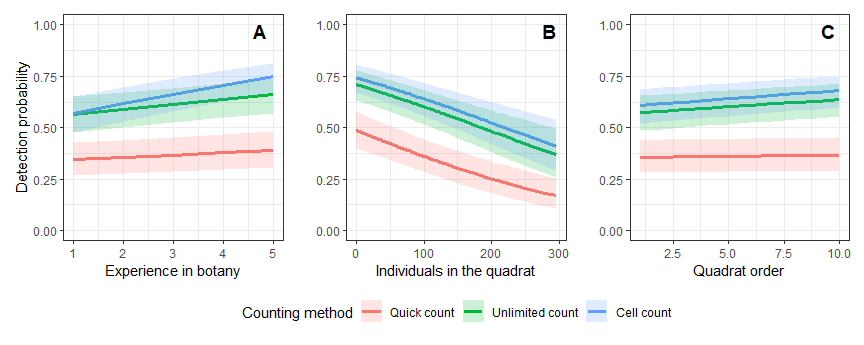

**Figure S7.3:** Predicted detection probability of individuals depending on the experience in botany of the observer (A), the true density (B) and the position of the quadrat in the round (C). The other variables included in the model are held constant at the following levels: species conspicuousness = 2, habitat closure = 0, quick count time = 0.5 min, unlimited count time = 3 min, cell count time = 3 min. Lines represent the mean detection probability of individuals, and the shading represents 95% confidence intervals.

### Appendix 8: Results with only the experienced botanists

The three figures below are the alternative versions of Figs 2–4 from the main text, made by fitting the model to a version of the dataset where only the observers with a high level of experience in botany were kept (i.e. levels of experience of 4 and 5 out of 5). This reduced the number of observations in the dataset from 4319 to 1070. This resulted in substantial differences in the parameter estimates and in the model’s predictions, but the general pattern of the predictions was unchanged. This indicates that our results are not due to an excess of observers with little or no field experience in our sample of observers.

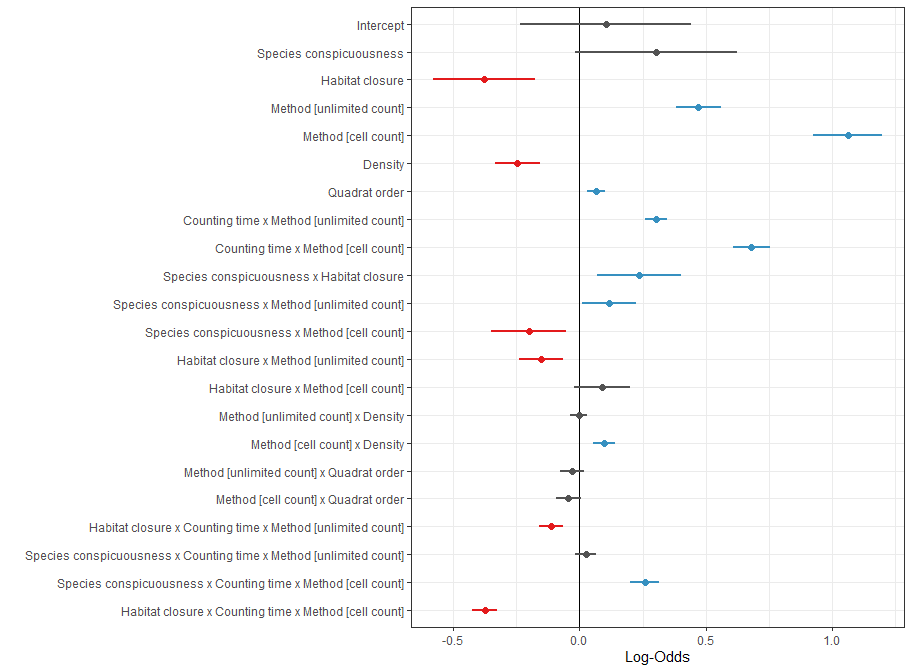

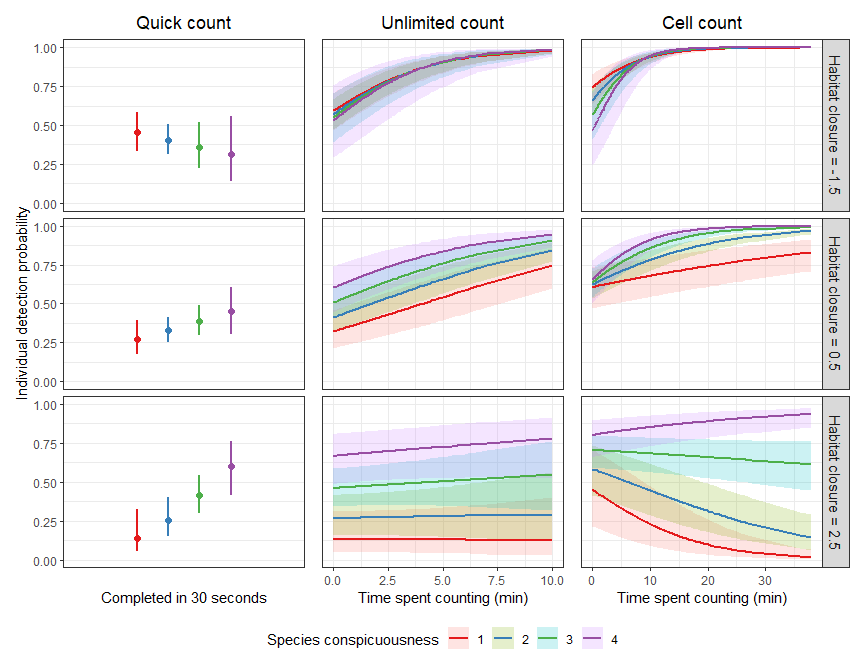

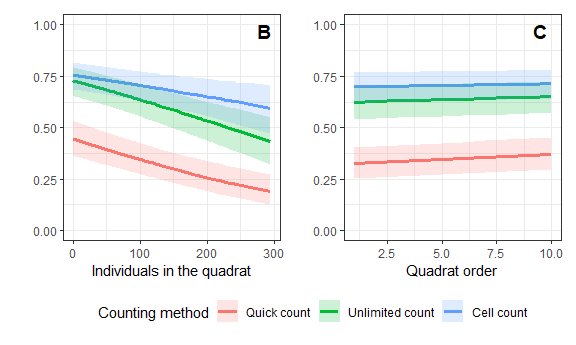

### Appendix 9: List of the participants

We would like to warmly thank the following 167 volunteers who participated in the experiment, some of them multiple times, including during the preliminary tests:

ALBERT Christophe, ALEGOT Bastien, AMARA Juliette, AMBROSIO DE LA IGLESIA Raquel, ANDRIEU Frederic, ARGAGNON Olivier, ARNAL Gerard, ARNAL Veronique, ASSIO Cindy, ASTRUC Guillelme, AUFFRAY Thomas, AUTHIER Louise, BARCZI Jean-Francois, BARRY Pierre, BASSIBEY Arnaud, BASTIEN Marion, BEAUTRU Lucas, BELGHALI Soumaya, BELLANGER Jean-Michel, BENVEGNEN Ulysse, BERNARD Charles-Etienne, BIOSSE Guilhem, BLANQUART Audrey, BLAYA Romane, BLIN Mirham, BLONDEL Francois, BONNET Lucie, BOULINIER Jenna, BOULY Ilona, BOUVIER Elodie, BRENDEL Frederic, BRIERE Maxime, CADET Serge, CANONNE Coline, CHAMBRELIN Justin, CHATELLIER Cyllene, CHAULIAC Christophe, CHAYRIGUES Sarah, CLOUET Louis, COCHENILLE Thomas, CONTOURNET Pascal, COULON Mireille, COUTURIER Thibaut, CUBAYNES Sarah, DARONAT Maelys, DE FRANCE Arthur, DECOMBEIX Anne-Laure, DECROCK Roxane, DELMOND Mathilde, DELPORTE Etienne, DEMONGEOT Marilou, DENIS Nans, DENTANT Cedric, DESPLANQUE Carole, DESPLAT Morgane, DIXON Lara, DORTEL Emmanuelle, DUBUISSON Candice, DUCRETTET Juliette, DUPUY Solene, DURRET Cassandra, ENGEL Julien, ESSELIN Mathilde, FALLOUR Delphine, FAUCHE Marine, FAURE Karine, FELIX Lila, FERRAILLE Thibaut, FERRER-LABROCHE Sophie, FICHTER Marc, FINOCCHIANO Marie, FLACHER Floriane, FONTAINE Ninon, FONTES Hugo, FORT Noemie, FORTUNY Xavier, FOUQUART Marilyne, FRAYSSE Remi, GANAULT Pierre, GAUTHIER Perrine, GENIEZ Philippe, GEOFFROY Sabine, GILBERTAS Lauriane, GILLIOT Marianne, GIRARDIER Marion, GRILLAS Patrick, GRITTI Clara, GUIRAUD Elise, HENRY Etienne, HOLVECK Pascal, HOPKINS David, HUYNH TAN Bernadette, ICARDO Emmanuel, IMBERT Eric, JANIN Romain, JEANNOT Anne, KACAMAK Begum, KANDEL Margot, KELLER Johann, KLESCZEWSKI Mario, LABROCHE Aurelien, LAQUEVILLE Manon, LARCHEY Enola, LARRE Antoine, LATRON Mathilde, LAUGIER Fanny, LAURET Valentin, LE BERRE Maelle, LE BORGNE Elsa, LEFRANCOIS Olivier, LLORENTE ZUBIRI Lucia, LOURENCO Marion, LUKAS Marie-Lou, MACE Bastien, MARQUES Fabien, MARQUIS Alois, MARRE Jacques, MARRE-CAST Josette, MASSART Pablo, MATUTINI Florence, MAUCLERT Virginie, MAURER Gilles, MEINERI Eric, MERTENS Louis, MEYER-BERTHAUD Brigitte, MIGAIROU Juliette, MOLINA James, MOLINO Jean-Francois, MUNZINGER Jerome, NACIRI Marwan, NADOT Sophie, PARIS Celia, PECHEUR David, PEROT Clara, PETIT Christophe, PIRES Mathias, POIRIER Clara, PONS Aurelia, PONS Virginie, POPOVITCH Yannick, POURTIER Laure, PRIEUR Jean, QUEDREUX Soham, ROCHER Leo, ROCHWERGER Naemie, ROLLIER Christophe, ROSSI Sofia, ROUYER Marie-Morgane, SAATKAMP Arne, SABY Lea, SAUVAJON Lou, SAVIO Laura, SEVIN Claire, SIMONNET Franck, TCHILINGUIRIAN Julien, TEMOIN Jean-Luc, TERREAU Alexandre, TESTUD Guillaume, TON Louis, TURPIN Louise, VELA Errol, VIDALLER Christel, VINCENT Gaelle, WASELLA Tatiana, YAVERCOVSKI Nicole, ZANELLA Agathe.
